# Stability of SARS-CoV-2 Phylogenies

**DOI:** 10.1101/2020.06.08.141127

**Authors:** Yatish Turakhia, Bryan Thornlow, Landen Gozashti, Angie S. Hinrichs, Jason D. Fernandes, David Haussler, Russell Corbett-Detig

**Author notes:** Equal contribution (Yatish Turakhia, Bryan Thornlow).

## Abstract

The SARS-CoV-2 pandemic has led to unprecedented, nearly real-time genetic tracing due to the rapid community sequencing response. Researchers immediately leveraged these data to infer the evolutionary relationships among viral samples and to study key biological questions, including whether host viral genome editing and recombination are features of SARS-CoV-2 evolution. This global sequencing effort is inherently decentralized and must rely on data collected by many labs using a wide variety of molecular and bioinformatic techniques. There is thus a strong possibility that systematic errors associated with lab-specific practices affect some sequences in the repositories. We find that some recurrent mutations in reported SARS-CoV-2 genome sequences have been observed predominantly or exclusively by single labs, co-localize with commonly used primer binding sites and are more likely to affect the protein coding sequences than other similarly recurrent mutations. We show that their inclusion can affect phylogenetic inference on scales relevant to local lineage tracing, and make it appear as though there has been an excess of recurrent mutation and/or recombination among viral lineages. We suggest how samples can be screened and problematic mutations removed. We also develop tools for comparing and visualizing differences among phylogenies and we show that consistent clade- and tree-based comparisons can be made between phylogenies produced by different groups. These will facilitate evolutionary inferences and comparisons among phylogenies produced for a wide array of purposes. Building on the SARS-CoV-2 Genome Browser at UCSC, we present a toolkit to compare, analyze and combine SARS-CoV-2 phylogenies, find and remove potential sequencing errors and establish a widely shared, stable clade structure for a more accurate scientific inference and discourse.

**Foreword:** We wish to thank all groups that responded rapidly by producing these invaluable and essential sequence data. Their contributions have enabled an unprecedented, lightning-fast process of scientific discovery---truly an incredible benefit for humanity and for the scientific community. We emphasize that most lab groups with whom we associate specific suspicious alleles are also those who have produced the most sequence data at a time when it was urgently needed. We commend their efforts. We have already contacted each group and many have updated their sequences. Our goal with this work is not to highlight potential errors, but to understand the impacts of these and other kinds of highly recurrent mutations so as to identify commonalities among the suspicious examples that can improve sequence quality and analysis going forward.

## Introduction

Extremely rapid whole genome sequencing has enabled nearly real-time tracing of the evolution of the SARS-CoV-2 pandemic [1–5]. By leveraging sequence data produced by labs throughout the world, researchers can trace transmission of the virus across human populations [6–14]. Typically, viral evolution is encapsulated by a phylogenetic tree relating all of the virus samples in a large set to one another [5,15–19]. However, despite efforts to mitigate the impact of sequencing and assembly errors, and to provide standardized datasets for real-time analysis [20], inferred phylogenetic histories of the outbreak often differ between analyses of different research groups (Results) and these inferred histories sometimes differ between analyses performed by the same group with different data (*e.g.,* 31 different Nextstrain trees produced between 3/23 and 4/30, Results). These differences may be created or accentuated when samples that contain unidentified sequencing errors are incorporated into the phylogenetic tree. Defining stable and easily referenced major clades of the virus is essential for epidemiological studies of viral population dynamics [17,18]. An understanding of how errors might be affecting the trees that are being published is essential to achieving that goal (Figure 1).

**Figure 1:**
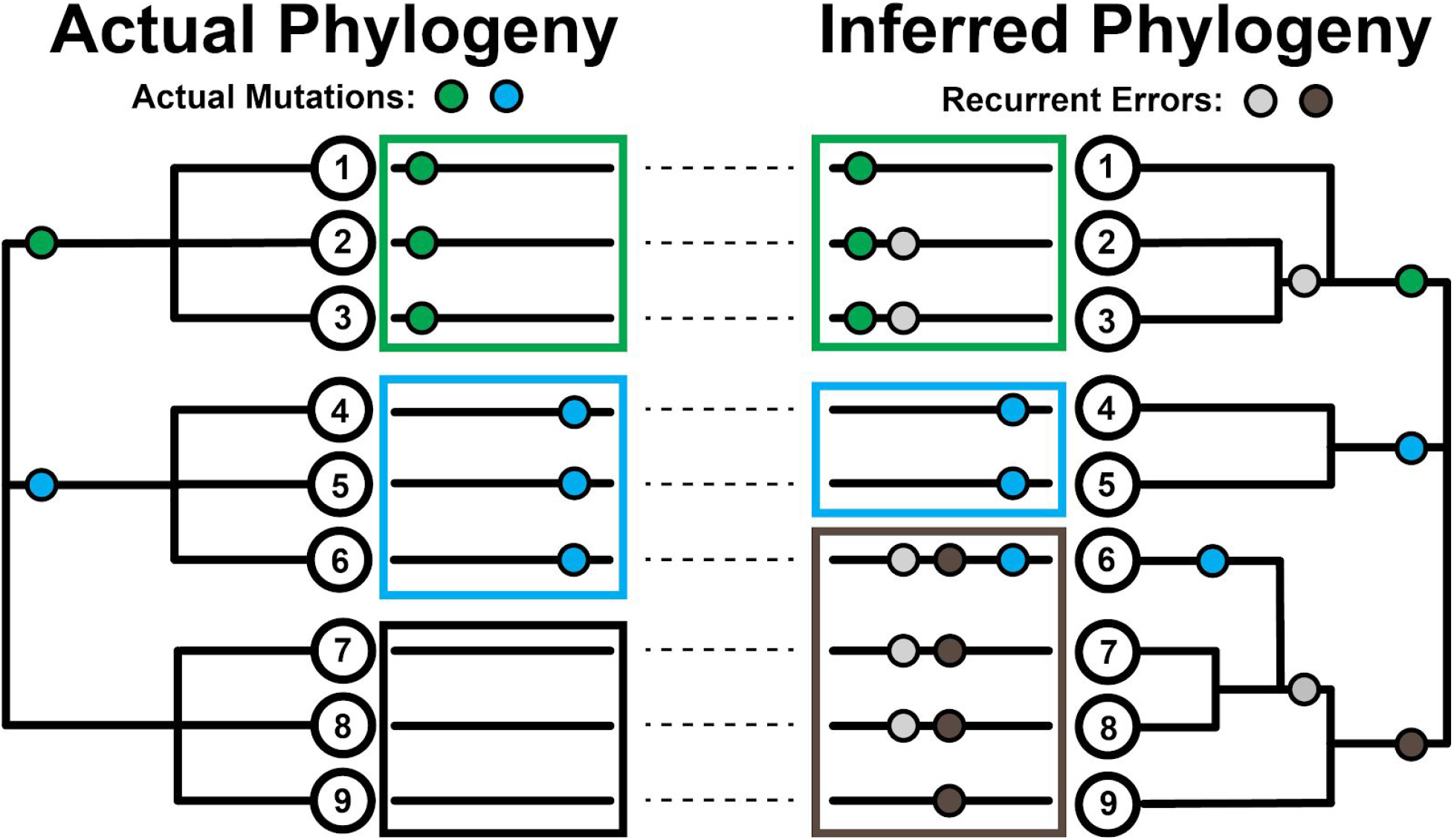
Effect of recurrent sequencing mutations on phylogenetic inferences. (Left) Pictorial representation of how the evolutionary histories of viral sequences (long black lines adjacent to tree nodes) can be traced on a phylogenetic tree using mutational events (green and blue circles) that occur. In this case, each mutation occurs once independently. (Right) The introduction of recurrent errors (gray and brown circles) can obscure the true evolutionary relationship between sequences leading to the creation of artifactual subgroups/clades (green-gray, leaves 2 & 3) and gray-brown, leaves 7 & 8)) and even the incorrect assignment of viral sequences to subgroups (leaf 6 no longer correctly groups with the blue subgroup (leaves 4 & 5); large boxes group together subgroups based on inferred first mutation).

It can be difficult to distinguish sequencing errors of different types from genuine transmitted and non-transmitted mutations in genome sequences. Taking a conservative approach, many researchers remove mutations that are observed only once during the evolution of the virus when constructing a phylogenetic tree, as these may be more likely to be errors [21,22], or non-transmitted mutations. However, systematic errors, where the same error from a common source is introduced many times in otherwise distinct viral genome sequences, are not removed by that approach [23,24]. These are more problematic, as they can appear as if they are genuine transmitted mutations (Figure 1). This might result from recurring errors in data generation or processing, or due to contamination among samples. Each case induces an apparent mutation that may be challenging to rectify with the real structure of the viral tree. Consequently, systematic errors can produce support for erroneous relationships between viral isolates and destabilize tree-building efforts. One possible approach is to mask out specific sites in the genome sequence where recurring errors are suspected, as suggested previously [24]. However, genuine recurrent mutations that may contain important information about properties of viral evolution [6,8,25–27] are sometimes hard to distinguish from recurrent systematic errors, and this could obscure important biology. Here, we present data that we hope will help the community make the important decision as to how to treat potential errors in SARS-CoV-2 genome sequences.

In addition to their influence on phylogenetic inference, recurrent systematic errors can also lead to erroneous inferences about viral mutation processes, recombination and selection. For example, artefactual biases in mutational processes could confound signatures of mutational hotspots [28–33]. The issue of whether or not recombination has occurred during the outbreak is critical to the immunological battle against the virus and is under intense debate [6,34–40]. Because many tests of recombination assume that all mutations can only occur once at each site, recurrent mutation and systematic errors can confound signatures of recombination [6,26,35]. Finally, recurrent mutations have been identified as a possible signatures of elevated mutation rates and natural selection in SARS-CoV-2 [8,13,24–26,29,33,35,41], but some of these apparent instances of selection may be due systematic errors in the sequences. Confusion about recurrent mutations and recombination affects our understanding of host response and influences our decisions about which viral molecular processes or specific immune epitopes we might want to target in vaccine development. Thus, it is essential that we explore the possible extent and impact of systematic errors in the viral genome sequences.

Another basic problem in current investigations of viral evolution is widespread phylogenetic uncertainty. In part, this has prevented the community from settling on a consensus definition of distinct viral clades (“(sub)types”, “groups”, “lineages”) representing the early divergence events, producing a “tower of Babel” problem in the scientific discourse [17,42]. Furthermore, many groups are making phylogenetic trees with widely varying goals, including dissecting patterns of nucleotide substitution, recurrent mutation, local lineage tracing, and large-scale phylogenomics [8,17,26]. The resulting topologies vary dramatically in structure, owing to differences in analysis choices and to phylogenetic uncertainty stemming from limited genetic diversity in the expanding viral populations. Consistent approaches for identifying commonalities and rectifying differences among trees are therefore foundational to the efforts to characterize viral evolution and epidemiology. A maximally stable topology will be essential for consistent nomenclature and facilitating conversations between analyses [17,42].

Our work takes on these two interrelated considerations: systematic errors and phylogenetic uncertainty. First, we show that hundreds of samples in the current SARS-CoV-2 sequencing datasets are affected by lab-associated mutations, which are potentially erroneous (see also [24]). These mutations distort phylogenetic inferences at scales most relevant to local lineage tracing and impact inferred patterns of mutational recurrence and recombination. We demonstrate that many can be identified and removed by cross-referencing patterns of recurrence against the source sequencing lab, and we provide automated methods for detecting suspicious and highly recurrent mutations. Second, to facilitate communication and comparison across different SARS-CoV-2 phylogenies, we develop approaches for efficiently comparing and visualizing differences among trees. All of the tools and functionality that we describe here are publicly available and integrated into the UCSC Genome Browser to facilitate rapid visualization, data exploration, and cross referencing among datasets and analyses. We anticipate that these methods will fuel more accurate continued discovery during the current pandemic and beyond.

## Results/Discussion

### Nextstrain Datasets

Our analyses are built in large part on the work of Nextstrain [15]. This team has already implemented a number of precautions to remove problematic sites and samples. In particular, they remove samples that are too divergent from others or whose date of sampling is inconsistent with the number of accumulated mutations. Additionally, all indels in the resulting multiple alignment are masked. Here, we do not consider the impact of alternative multiple alignments, upstream filtering methods, or the possible impacts of indels. Each of these factors has the potential to affect downstream analyses and should be considered carefully. For our purposes, we anticipate that Nextstrain’s filters will minimize idiosyncratic errors and should be retained in the majority of future analyses. Here, we use as a primary example, 31 different Nextstrain trees from days between 3/23/2020 and 4/30/2020. We focused in particular on the dataset from 4/19/2020 which contains 3246 variants in total (Methods). The vast majority of variants are at low frequency, as is expected for a rapidly expanding population.

### Systematic Error Could Be Mistaken for Recurrent Mutation or Recombination

Non-random errors can present a fundamental challenge for phylogenetic inference and to the interpretation of viral evolutionary dynamics. There are at least four possible sources of (real or apparent) mutations that recur within independent lineages in a tree, and each makes distinct predictions about the source of recurrent mutations (Table 1). In particular, recent work has shown a strong bias towards C>U mutation in the SARS-CoV-2 genome [21,42–44]. Systematic errors, which usually result from consistent errors in molecular biology techniques or bioinformatic data data processing, need not reflect this bias and are not subject to natural selection. We therefore anticipate that many systematic errors will affect many mutation types, modify protein sequences, and strongly correlate with genome sequences generated in particular labs [24].

**Table 1.** Expectations for each source of apparent recurrent mutation.

| Source | Heritable | Typical Allele Frequency | C>U Biased | Mutation Bias? Minor vs major | Lab Correlation | Extremal |
| --- | --- | --- | --- | --- | --- | --- |
| Recurrent Mutation | Y | Low | Y | Y | N | Y |
| Recombination | Y | High | Y | N | N | N |
| Systematic Error | N | Low | N | Y | Y | Y |
| Contamination Error | N | High | Y | N | Possible | N |

### Many Apparently Recurrent Mutations Found in the SARS-CoV-2 Genome

To examine patterns of recurrent mutation we employ a simple statistic, the parsimony score, which is the count of the minimum number of unique mutation events consistent with a tree and sample genotypes ([45,46] computed using our software from https://github.com/yatisht/strain_phylogenetics, Methods). More sophisticated statistics could be employed, but this simple one is effective, is readily interpretable, and can be computed rapidly. We restrict most analysis to bi-allelic sites, i.e. sites that contain one the allele in the reference genome from the root of the tree (here and in Nextstrain this is, Wuhan-Hu-1, obtained in December 2019 in the city of Wuhan) and a single alternate allele. Across the 4/19/2020 Nextstrain tree, we found 2533, 395, 94, 40, and 44 bi-allelic sites with parsimony score one, two, three, four, and five or more, respectively (Figure 2, Figure S1). In particular, there is a strong “on diagonal” component of the data that is defined by a linear relationship between the log of the alternate allele count and parsimony score (dashed line in Figure 2A, log2-based slope = 3.188). These mutations reoccur across the phylogeny at exceptional rates relative to their allele frequencies. Hereafter, we refer to the set of variants in this on-diagonal group as **extremal** sites (blue, red, and orange in Figure 2A). This relationship suggests that the extreme accumulation of independent clades for the alternate allele is logarithmically related to the number of instances of the alternate allele in the phylogeny (Figure 2B). This suggests that even the most mutable or error prone sites in the genome will sometimes have alternate alleles grouped into clades during phylogenetic inference thereby appearing to be inherited.

**Figure 2.**
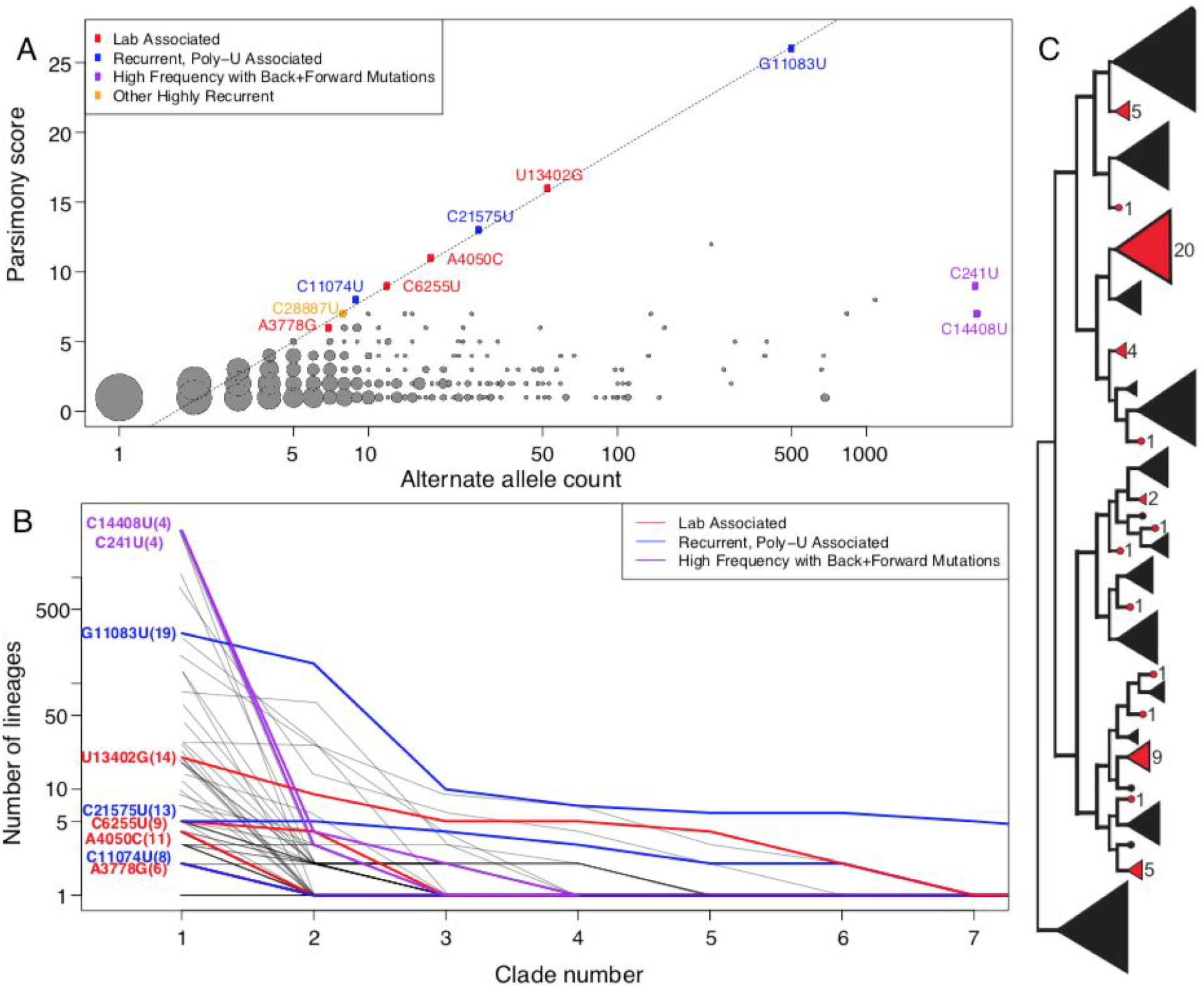
**(A)** The relationship between alternate allele count and parsimony score. Point radius indicates how many sites share a single parsimony score and alternate allele count. Several noteworthy recurrent mutations are labelled. Note that the X-axis is log-scaled. **(B)** The sizes of independent clades for the same alternate allele arranged in descending order. The number of lineages per clade is shown on logarithmic scale facilitating comparison with Panel (A). These data indicate that when alternate allele clade sizes for a given site are sorted in decreasing order, their sizes are reduced going from left to right by a multiplicative factor at each step, consistent with the log-linear relationship displayed in Panel (A). Mutations with remarkably high recurrence are shown with color reflecting their properties: lab-associated (red), recurrent and associated with a poly-U stretch (blue), and high frequency with many forward and backward mutations (purple). Grey lines in the background are the same values but for all other mutations with parsimony score 4 or greater. The values in parentheses in the mutation names indicate the number of unique clades associated with the alternate allele. Note that in some cases, this extends beyond the limit of the X-axis and that the Y-axis is log-scaled for visibility. **(C)** An example of the observed patterns of evolution at one highly recurrent site with reference allele U and alternate allele G, site 13402 and parsimony score 14, where 14 alternate allele clades (in red) each represent an apparently independent incidence of the mutation substituting the alternate allele.

### Automated Detection of Extremal Sites

Lab-association is a straightforward indication that we use below to identify highly suspect mutations in the SARS-CoV-2 genome. However, hypermutable sites might also impact phylogenetic reconstruction for similar reasons as systematic errors. We therefore sought to provide researchers with a method for rapidly identifying and flagging suspiciously recurrent mutations. We therefore developed code to identify the “on diagonal” extremal sites and produce plots of the output similar to Figure 2, that is available at https://github.com/yatisht/strain_phylogenetics. Note that depending on the dataset, this component is not always so linear as in Figure 2, but it is associated with highly homoplastic sites regardless (*e.g.,* Figure S1). Our list of extremal sites includes two that we later show are strongly lab-associated (A4050C and U13402G), three mutations that are adjacent to >5bp poly-U segments in the genome (C11074U, G11083U, and C21575U), as well as two more C>U mutations (C21711U, C28887U). Regardless of their proximate causes, highly recurrent mutations can negatively impact the accuracy of inferred tree topologies, and thus should be removed prior to phylogenetic tree construction and for many subsequent analyses.

### SARS-CoV-2 Data Contains Many Lab-Associated Mutations

To search for systematic errors associated with a particular lab, we extracted the set of sites with parsimony score 4 or more. We then flagged sites as lab-associated mutations if more than 80% of the samples containing the alternate allele were generated by a single group. Using this heuristic approach, we found 16 such sites (Table S1). We note that this set of sites contains two mutations previously identified as lab-associated mutations [24], some others identified as highly homoplastic [8,24,25,42], as well as several identified as evidence for recombination [26]. These mutations in lab-associated sites display a range of base compositions and only one is a C>U transition (C6255U). This rate of C>U mutation is much less than the genome-wide average rate of C>U mutation for non-singleton sites (49%, P = 0.0004914, Fisher’s exact test), and differs significantly from the rate of C>U mutation among our set of highly recurrent mutations that are not strongly associated with a single sequencing lab (P = 1.005e-07, Fisher’s exact test). Furthermore, our set of lab-associated mutations is weakly enriched for protein altering mutations relative to other highly recurrent mutations (P = 0.09372). Collectively, our results suggest that some recurrent mutations among these 16 could be lab-associated systematic errors.

The potential causes of lab-associated mutations are numerous. A non-exhaustive list follows. First, primers for reverse transcription or PCR might introduce systematic errors either via errant priming, because they “overwrite” true variation, or because of errors during bioinformatic processing. For example, the commonly used ARTIC primer sets amplify the viral genome from metatranscriptomic cDNA by tiling the viral genome with PCR amplicons (https://artic.network/). Second, if a portion (perhaps a single amplicon) from a contaminating sample were present in many sequencing reactions from a single lab, this could propagate variants across all genome sequences from a single group. Third, contamination from the human transcriptome itself might be inadvertently included in assembled viral genomes.

Two labs contributed a disproportionate number of lab-associated mutations in our dataset, suggesting a consistent source of these alternate alleles (Table S1). One lab group is strongly associated with two adjacent high parsimony score mutations A24389C and G24390C. These occur in a 10bp sequence that otherwise closely resembles an Oxford Nanopore sequencing adapter, CAGCA**CC**TT, and is adjacent to an ARTIC primer binding site. Here, the differences between the genome sequence and adapter are bolded. See also [24], where a commenter on that work comes to a similar conclusion regarding the likely source of these mutations. Additionally, A4050C, U8022G, U13402G, and A13947U (Figure 2, Table S1) are associated with this same lab and either overlap or are within 10bp of ARTIC primer binding sites (14_left_alt4, 26_right, 44_right, and 47_left, respectively), suggesting that a consistent bioinformatics data processing error may be responsible. Sequences submitted by another lab group are strongly associated with four additional high parsimony score mutations, G2198A, G3145U, A3778G, and C6255U (Figure 2, Table S1). Here again, each of these intersects one of the ARTIC primer binding sites (8_left, 11_left, 13_left and 20_right respectively, Figure 3). In aggregate, our set of lab-associated mutations are significantly closer to ARTIC primer binding sites than would be expected by chance (P = 0.0283, permutation test, Figure 3). Our results therefore suggest that mutations intersecting or immediately surrounding commonly used primer binding sites should be subjected to particular scrutiny.

**Figure 3.**
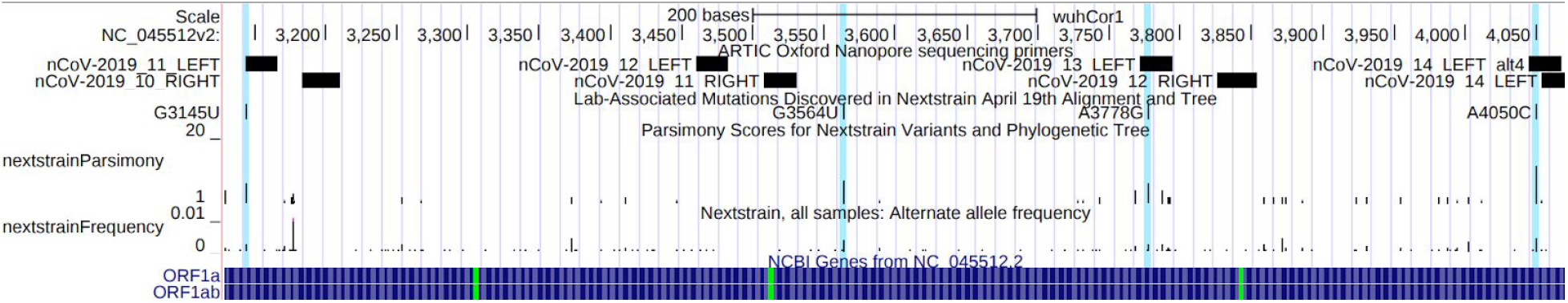
UCSC Genome Browser display of lab-associated mutations and ARTIC primers. Bases 3130 to 4070 of the SARS-CoV-2 genome are displayed, containing four lab-associated mutations highlighted in light blue. G3145U, A3778G and A4050C overlap ARTIC primer bind sites. An interactive view of this figure is available from: http://genome.ucsc.edu/s/SARS_CoV2/labAssocMuts

Another lab-associated mutation, C22802G, also overlaps an ARTIC primer (76_left, Table S1), but the ultimate source is unrelated. In this case, that would not be possible because these SARS-CoV-2 genomes were assembled from whole metatranscriptomic data without PCR selection. Instead, the cause appears to be misalignment of a human ribosomal RNA sequence that was incorporated into the consensuses for a subset of genomes produced by this group (Dr. Darrin Lemmer, *Pers. Comm.*). This highlights the broad range of possible causes of lab-associated mutations.

It is more challenging to identify the specific sources of the other five lab-associated mutations that we observed, but commonalities are informative. Three of these mutations are associated with a single group and each is a G>U transversion (G3564U, G8790U, G24933U, Table S1). Even more strikingly, each mutation occurs in a GGU motif, suggesting a common molecular mechanism might underlie this set of lab-associated mutations as well (*i.e.,* GGU > GUU). One possible hint is that this group uses a transposase-based library preparation method, which is relatively uncommon among SARS-CoV-2 sequencing groups and might explain this unique signature. Beyond these, G1149U and U153G are associated with two different sequencing groups, but do not show similar signatures as other variants (Table S1). More generally, the fact that many recurrent mutations are associated with genome sequences produced by individual lab groups suggests that consistent data processing or generation issues affect many sites. For example, sample contamination, which can be quite challenging to confidently detect, might also contribute to mutational recurrence and might not strongly be lab-associated (Text S1). However, we caution that this does not definitively prove that these apparent mutations are errors, but we believe it is prudent to remove these sites for most analyses until additional sequencing corroborates them.

### Lab-Associated Mutations are Consistent with Simulated Systematic Error

To study how systematic errors affect phylogenetic inference and inferred properties of viral evolution, we experimentally introduced errors in replicate experiments. We found that the parsimony score displays a roughly linear relationship with the log of the alternate allele count, as it does for extremal sites in Nextstrain trees we examined built on different days in April, but with varying slope (Figure 4). This is expected because errors will sometimes occur in sample genomes whose positions are close on the real phylogeny and even in sister lineages. Tree-building methods could then group these samples into a single clade. Importantly, the effect of drawing samples together can cause systematic error, or hypermutable sites for that matter, to appear heritable.

**Figure 4:**
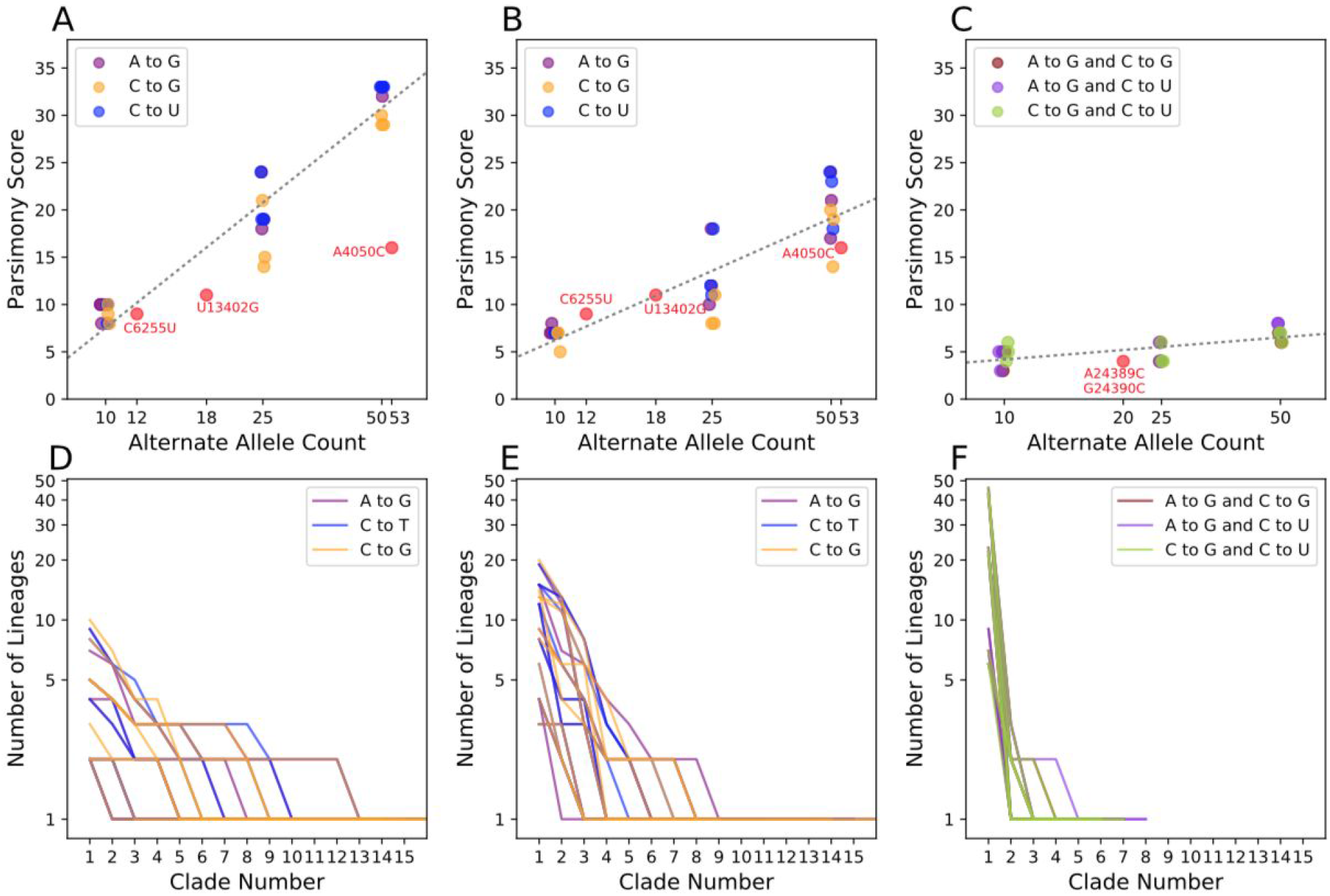
Parsimony scores at sites with introduced systematic errors. We added artificial errors to 10, 25, and 50 Australian (A) and early-March French (B) samples at the sites A11991G (purple), C22214G (blue), and C10029U (orange) in three replicates, then produced phylogenies and computed the parsimony score at each site. (C) We also introduced errors to the early-March French samples two at a time per sequence rather than individually. For comparison, we also show the values for three lab-associated mutations (C6255U, U13402G, A4050C; A, B) and for pair of linked lab-associated mutations (A24389C and G24390C; C). Each panel (A-C) contains a best-fit line (as in Figure 2A), for the relationship between log2 alternate allele count and parsimony in simulated error data (slopes = 10.0, 5.55, and 1.0). (D-F) Corresponding clade sizes arranged in descending order for error simulations in (A-C, respectively, as in Figure 2B).

Additionally, we find that viral genetic background and mutation type is an important contributor to this relationship. When errors are placed randomly across Australian samples (Figure 4A), we see much higher parsimony scores than when errors are placed only in samples from France collected between March 1 and March 17 (Figure 4B). The difference likely reflects the fact that the samples from France are more closely related. Because many of the lab-associated mutations that we identified are derived from a similarly restricted time and geographic region as our samples from France, parsimony scores at those sites closely resemble these sets of simulated error (Figure 4B). This suggests that the identification of lab-associated mutations will become increasingly straightforward as the viral populations accumulate genetic diversity. We also observe that mutations that truly occur less often during SARS-CoV-2 evolution (*e.g.,* C to G) have slightly lower parsimony scores. This is likely due to modelling nucleotide-specific mutation rates during tree-building where mutations consistent with viral mutational processes are less likely to be erroneously grouped. Importantly, our results suggest that a simple heuristic based on each site’s parsimony score and recurrence is sufficient to identify most lab-associated mutations above very low frequencies. However, extremely infrequent lab-associated error could be challenging to distinguish from more conventional sequencing error.

Because systematic errors also affect the inferred tree, they can impact inferred patterns of mutational recurrence at other positions in the genome as well. In 50 out of 54 total experiments where we introduced a single recurrent error, we found that the parsimony score increased at other sites (range 2 to 44). This emphasizes the importance of identifying and excluding such mutations prior to inferring the final tree and downstream analyses.

### Correlated Lab-Associated Mutations Have Large Impacts on Phylogenetic Inference

If infrequent but highly correlated errors were introduced at different sites in many samples, this could cause more samples to be grouped into a clade. We might not easily detect these errors based on recurrence. Two lab-associated mutations, A24389C and G24390C, are not just on adjacent genomic locations but are nearly perfectly correlated across samples. These sites have low parsimony scores when compared to other lab-associated mutations (4 and 5, respectively, Figure 4C). When we introduced similar correlated errors, we found that the parsimony scores were lower than in single error introduction experiments. Nonetheless, in only two error introduction experiments (out of 9) with 10 affected samples did we see a parsimony score as low as 3. Although low frequency and highly correlated error could be challenging to identify in general, we believe this is infrequent in our dataset (see Text S2). Therefore we have not included tests for correlated errors in our suggested methods for finding lab-associated mutations, but adjacent correlated sites should be carefully scrutinized.

### Lab-Associated Mutations Affect Phylogenetic Inferences on Scales Relevant to Local Lineage Tracing

To investigate the impacts of lab-specific mutations on phylogenetic inference, we removed (“masked”) each of the 16 sites with a lab-associated mutation (Table S1). Importantly, removing lab-associated mutations sometimes impacted phylogenetic patterns at other sites. For example, after removing all lab-associated mutations, the evidence for back-mutations at C14408U is eliminated, while many forward-mutations remain (*e.g.*, Figure 5). In fact, the parsimony score changed for 107 sites and decreased for 53 sites on the tree that we inferred after removing all of the lab-associated mutations relative to the tree inferred including all sites. Additionally, we find that many samples containing lab-associated mutations have been repositioned on local topologies (*e.g.,* Figure 5). Furthermore, in some cases the placement of closely related lineages that are unaffected by lab-associated mutations is also affected (Figure S2). These mutations therefore affect phylogenetic inferences at scales relevant to local lineage tracing, which may obscure dynamics of local transmission.

**Figure 5.**
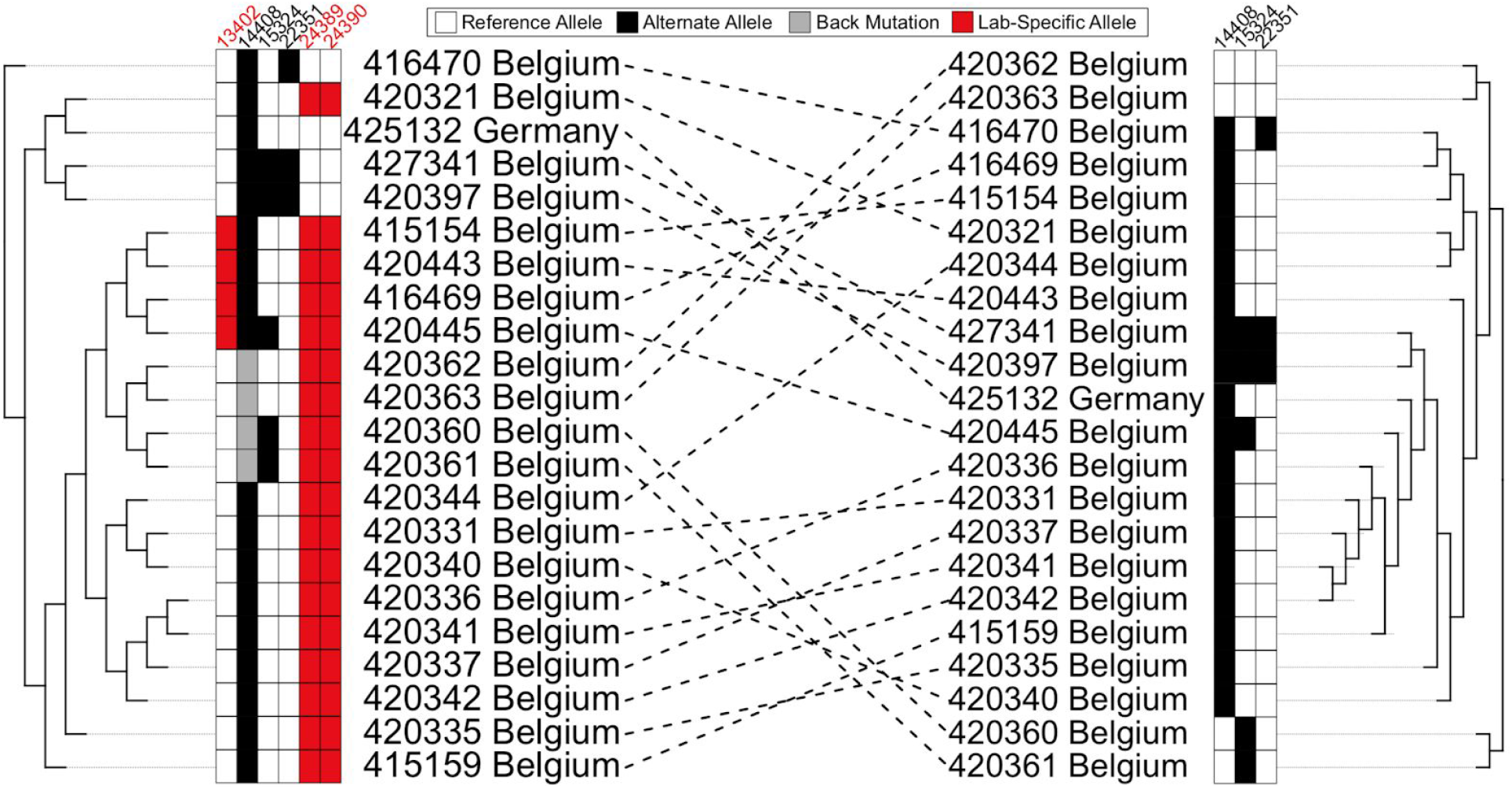
Lab-associated mutations impact phylogenetic inferences. Part of the tree we inferred from the 4/19/2020 Nextstrain dataset (left) compared to the corresponding part of tree after removal of sites with lab-associated mutation (right). Lab-associated mutations (red) can affect the inferred phylogeny and are associated with apparent back-mutation to the ancestral allele (grey in column 14408, left) at other sites (white). When lab-associated mutations are removed, the resulting tree (right) shows no evidence for back-mutation at those sites (now white in column 14408), though several independent forward mutations remain evident.

To examine the effect of each lab-associated mutation and the other extremal sites in isolation from one another, we individually masked each site and inferred a phylogeny. As a comparison, we also masked a set of sites that have similar alternate allele frequencies as the lab-associated mutations, but each has a parsimony score of one. The distributions of entropy-weighted total distance (a measure of distance between trees, described below) are remarkably similar when masking individual lab-associated sites, other extremal sites, and our control sites (Figure 6). Most exceed the distance we observed when we independently inferred two trees from the same input alignment (dashed black line). Our results therefore suggest that the lab-associated and extremal sites can impact tree-building approaches on par with real mutations, although the effects are typically small on the scale of whole topologies, as is expected given their typically low allele frequencies (Figure 6, Figure S3).

**Figure 6:**
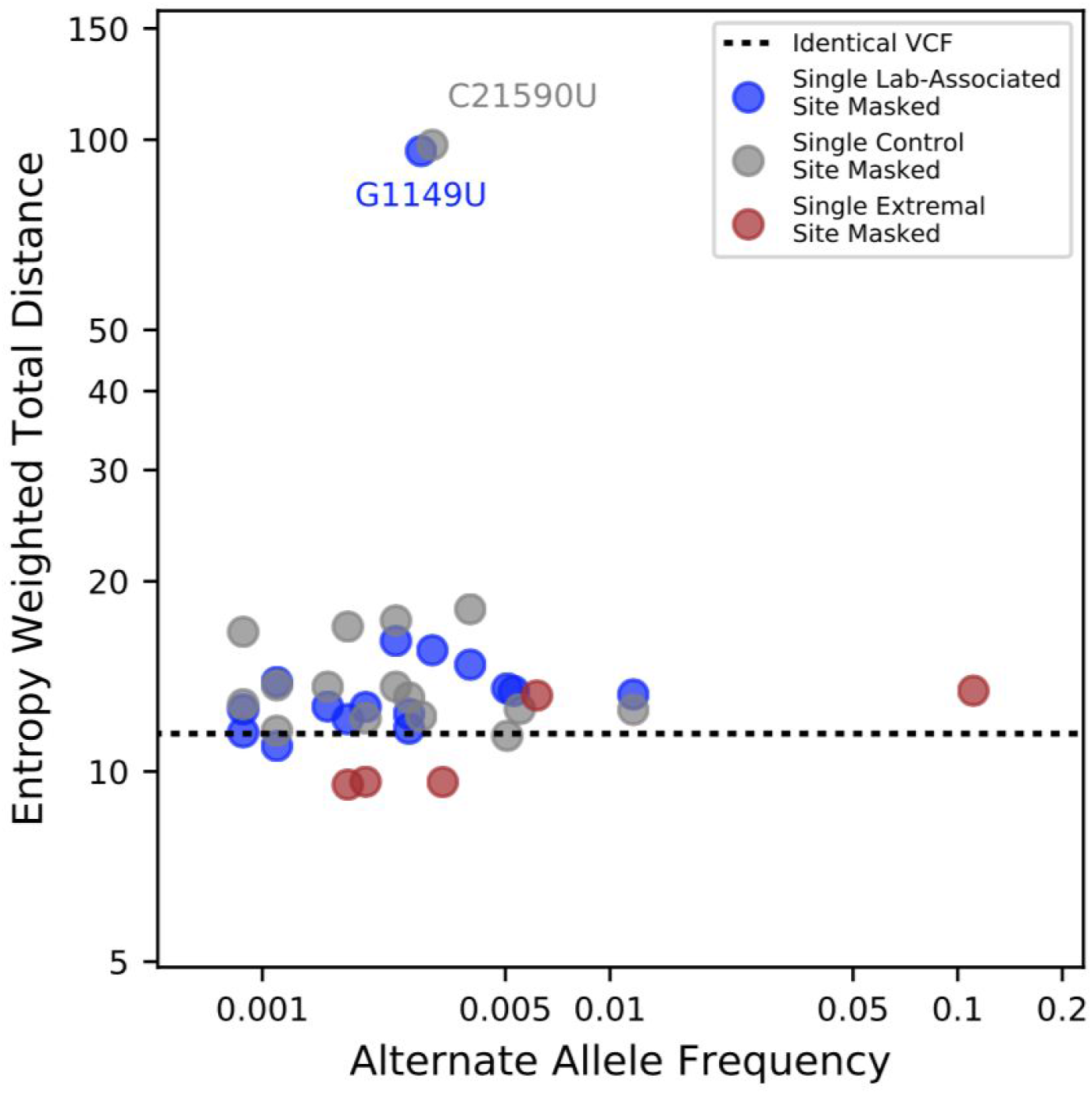
The relationship between alternate allele frequencies of lab-associated mutations and effect of masking on inferred tree topology. Entropy-weighted total distances relative to the reference maximum likelihood phylogeny are shown for phylogenies constructed after masking individual sites. Blue points correspond to sites with lab-specific alternate alleles, grey points correspond to control sites with parsimony scores of 1 and similar alternate allele frequencies to the sites with lab-specific alternate alleles, and brown points correspond to non-lab-specific extremal sites. The black horizontal line indicates the entropy-weighted total distance value for a maximum likelihood phylogeny constructed from an alignment identical to that of the reference phylogeny. Two outliers, C21590U (control) and G1149U (lab-associated) have outsize effects on inferred tree topology.

Phylogenies made after removing two mutations, one control and one lab-associated are outliers for entropy-weighted total distance (Figure 6, Figure S4) and other tree distance statistics (Figure S3). In each case, however, the likelihood of the tree produced from the full dataset is actually higher (Table S5), suggesting that our tree-building method discovered a different locally optimal but less favorable topology rather than a dramatic impact of each site individually. These results suggest higher level uncertainty in the tree topology largely independent of the effects of lab-associated mutations.

### Recurrent Mutations Not Associated with a Lab Reflect the Mutation Spectrum of The SARS-CoV-2 Genome

Hypermutation rather than positive selection may explain many remaining highly recurrent sites. Previous analyses showed that the rate of C>U mutation is exceptionally high relative to other mutation types in the viral genome [10,25,43,47]. This class of mutations should show increased evidence of recurring multiple times because they experience elevated mutation rates [25]. Indeed, parsimony scores at sites containing C>U mutations are significantly higher than those for all other mutation types (P < 2.2e-16, Wilcoxon Test, Figure 7). Furthermore, parsimony scores at C>U sites also significantly exceed those at G>A (P = 5.993e-12) as well as U>C (P = 1.407e-10) sites. This mutational bias might be driven by APOBEC editing of the viral genome [25,43,44,47–49]. Consistent with previous results [43,44,47–49], we find that 5’-[U|A]C>U mutation more frequently than 5’-[C|G]C>U (P = 0.0501), but we do not see a similar effect for 3’ flanking sites at 5’-C>U[U|A] relative to 5’-C>U[G|C] mutations (P = 0.378). The highly biased spectrum of C>U mutations and the correlation with local sequence context implies that the plus-stranded virus biology may be leading to recurrent C>U mutations [47].

**Figure 7.**
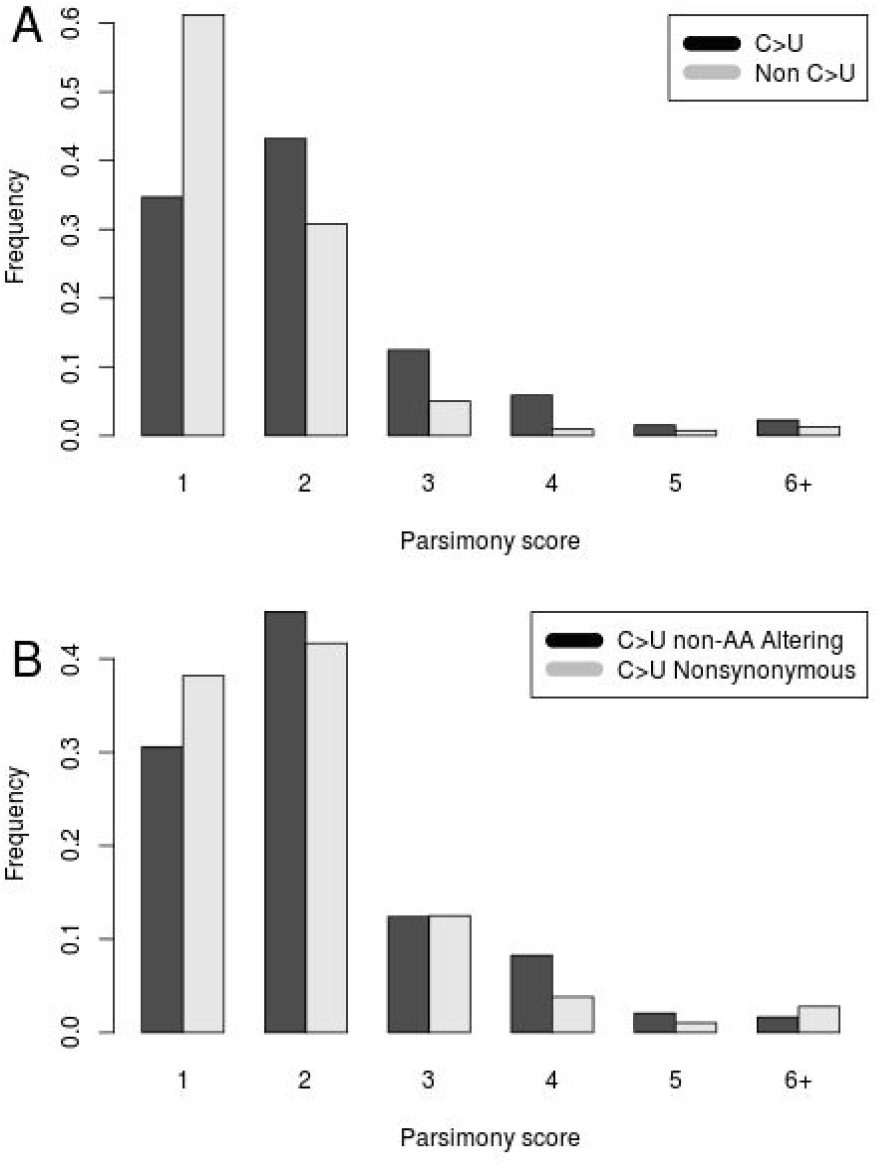
Recurrence of mutations during SARS-CoV-2 evolution. **(A)** Frequencies of parsimony scores for C>U (Black) vs all other mutation types (Grey). **(B)** Frequencies of parsimony scores for C>U mutations that do affect amino acid sequences (Grey), and those that do not affect amino acid sequences (Black).

Of the 83 highly recurrent mutations with parsimony greater than three, 50 are bi-allelic, not strongly lab-associated, and have an alternate allele frequency less than 0.01. Of these, 42 are C>U mutations. This is a significant excess of C>U mutations relative to the rate among non-singleton bi-allelic sites with parsimony score three or fewer (P = 3.658e-07, Fisher’s exact test). Additionally, C>U mutations that do not affect the underlying amino acid sequences display higher parsimony scores than do C>U mutations that do affect amino acid sequences (P = 0.0553, Wilcoxon test, Figure 7). This suggests that negative selection has played a role in shaping the distribution of highly recurrent mutations by purging strongly deleterious alleles.

Evidence suggests that any contribution of sequencing error to the excess of C>U mutation is small. Alternate alleles at 81.4% of sites with parsimony greater than 3 are corroborated by more than one sequencing technology. Of those, 77% of bi-allelic sites are C>U transitions (Table S2). Illumina C>T errors in raw sequence reads are typically enriched in the contexts of flanking G regions [50,51], but here we do not see this pattern. Similarly, nanopore sequencing typically creates errors in homopolymer stretches [52], but we only see a few recurrent mutations associated with such regions. Notable exceptions are the extremal sites C11074U, and C21575U, which abut poly-U stretches in the genome and might result from replication slippage (see also G11083U). It is possible that the excess of C>U mutations are driven in part by high error rates during reverse transcription [53–56], which is required for cDNA sequencing. However, C>U mutation is overrepresented in high frequency mutations as well (9/20 frequency > 0.025 mutations are C>U, Table S3), indicating that this bias likely reflects a true mutational process. Additionally, these mutations are approximately as distant from ARTIC primer binding sites as we would expect by chance (P = 0.7851, Permutation Test, Table S2). Collectively, our results suggest that neither library preparation or sequencing error is not the major driving force behind biased C>U mutation observed at highly recurrent mutations that are not strongly associated with a single lab. However, even if real, the existence of these highly recurrent mutations does not require that they are heritable (it must be that many viral mutations are never transmitted), in which case their phylogenetic behavior should be the same as systematic errors.

### Possible Mitigations for Lab-Associated and other Highly Recurrent Mutations

We proposed a simple heuristic approach to detect lab-associated mutations. To briefly reiterate here, first we identify sites that experience mutations on at least four independent branches of the SARS-CoV-2 tree, and then we extract the set where 80% or more of the alternate allele comes from sequences produced by a single lab. These are classified as lab-associated recurrent mutations. Then for all sites we plot parsimony score versus log2 of alternate allele count and determine a set of extremal sites as described in Methods. We recommend that lab-associated and most extremal mutations be masked for the purposes of constructing a phylogenetic tree to be used in downstream analyses. One exception here is extremal site 11083, which is sufficiently high frequency that it affects inference of the deepest branches of the tree. We suggest that it should be included during phylogenetic inference. However, alternative masking strategies that remove small clades containing apparent forward and backwards mutations at high frequency sites might also be effective and will be investigated going forward. Many downstream analyses following tree-building should consider masking 11083 as well. After masking the set of lab-associated and extremal sites, the samples which previously contained them can be retained in phylogenetic inference and downstream analyses. Tracks identifying these sites are available on the UCSC Genome Browser and in Table S1.

Though not a focus here, we emphasize that filtering for genomic regions that are difficult to assemble or align (*e.g.,* those used by Nextstrain to filter the ends of chromosomes as defined here https://github.com/nextstrain/ncov) should also be rigorously employed. In fact, in light of our discovery of a possible lab-associated mutation at position 153, which is just within the usual filtering range, we suggest that it may be preferable to simply mask the full 5’ and 3’ UTR regions, which are typically harder to assemble and align confidently.

To examine the aggregate effect of lab-associated and extremal mutations, we inferred a tree for the full dataset, and another after masking all lab-associated and extremal mutations except 11083 using IQ-TREE 2 with 1000 ultrafast bootstraps [57,58]. We then collapsed all branches that do not contain a mutation into a polytomy. In contrast to the single site masking experiments above, here the topologies of the two maximum likelihood consensus trees differ significantly. The symmetric entropy-weighted total distance between the two topologies is not large, 9.4, but the fit to the multiple alignment having masked these sites improved by 189 log-likelihood units relative to the tree inferred without lab-associated and extremal mutations. Below, we show that confident relationships at higher branches in the topology are minimally affected relative to other widely-used phylogenies, which were inferred including lab-associated and extremal mutations. Our phylogeny produced following these masking recommendations is available from the UCSC Genome Browser (Figure 8), and we will update and maintain this resource as we add new data, as other suspicious mutations are identified, and as improved masking recommendations are developed.

**Figure 8.**
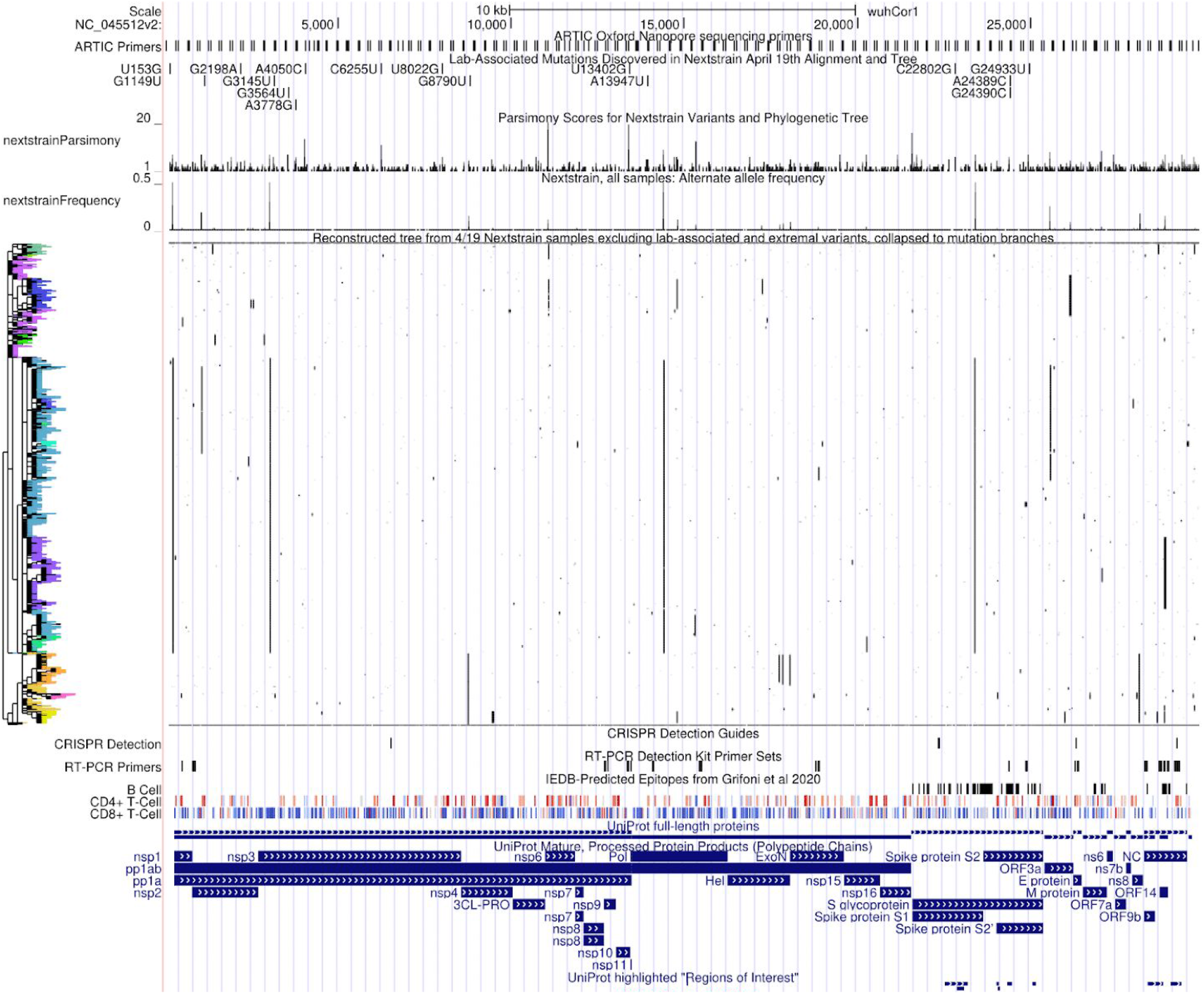
UCSC Genome Browser view of all lab-associated mutations in the context of parsimony scores, alternate allele frequencies, the full genetic variation dataset with phylogenetic tree constructed after removing lab-associated and extremal mutations. This genetic variation data can be cross-referenced against many other diverse datasets available in the UCSC SARS-CoV-2 Genome Browser. Interactive view: http://genome.ucsc.edu/s/SARS_CoV2/labAssocMutsAll

Many of the most intriguing and evolutionarily relevant biological phenomena, such as viral recombination and recurrent mutation, explicitly require inferences based on homoplastic mutations. Special caution is clearly warranted. For these analyses, it is still necessary to mask lab-associated mutations and extremal sites because they can destabilize phylogenetic inference, but clearly one could not exclude all homoplasies. In light of significant phylogenetic uncertainty, which we address below, we recommend that each analysis be repeated across alternative possible tree topologies to confirm the robustness of biological inferences. However, this is not without significant challenges and the most general solution for confirming recurrent mutation or recombination is heritability. If a mutant or recombinant lineage grows sufficiently large and is corroborated by many labs, we can be much more confident [26]. We therefore suggest that evidence of heritability and independent sequence confirmation should be required to support inferences of either recurrent mutation or recombination.

### Exploring Data Quality and Mutational Recurrence Using Our Tools

SARS-CoV-2 sequence data is growing at an incredible pace. Here we developed tools to enable investigations of similar patterns in updated and additional datasets. To summarize: (1) we provide a method for rapidly computing parsimony scores to identify highly recurrent positions; (2) we provide an approach for identification of unusually recurrent sites relative to their allele frequencies (here, termed extremal); (3) we provide an approach for semi-automated metadata correction (See Methods), which improved detection of lab-associated mutations; and (4) we provide a method for identifying the set of highly homoplastic mutations that are strongly associated with individual sequencing labs. Our heuristic cutoffs appear to perform well in the datasets we examined, but the program is designed to empower users to explore other datasets and other filters as well. Software to perform each analysis are provided via GitHub (https://github.com/lgozasht/COVID-19-Lab-Specific-Bias-Filter and https://github.com/yatisht/strain_phylogenetics).

### Visualizing Data Quality, Genetic Variation and Correlation via the UCSC SARS-CoV-2 Genome Browser

Data visualization remains one of the most powerful mechanisms for identifying unusual patterns and possible errors in genome sequence data (*e.g.,* Figure 3, above). Therefore, as an integral part of this work, we provide powerful data exploration and visualization tools that can be applied to future variation datasets as well. Output from our programs for computing parsimony scores and detecting lab-association mutations can be imported directly into the UCSC SARS-CoV-2 Genome Browser [59] as custom tracks to facilitate visual exploration of suspect mutations with a user defined vcf file and tree. This is a very useful visualization framework for data quality control and for investigating the root causes of highly recurrent mutation. For example, it is straightforward to explore the relationships between phylogeny, genetic variation, and functional genomic annotations (Figure 8).

Researchers can upload their own aligned SARS-CoV-2 genome samples and phylogenetic trees to the SARS-CoV-2 Genome Browser in order to compare their phylogenetic analysis to those from Nextstrain and COG-UK, and also to look at the specific molecular features of the clades that their phylogenetic analysis identifies (Text S4). These molecular features include widely used primer pairs (as in Figure 3) as well as CRISPR guides, predicted and validated epitopes for CD4+ and CD8+ T-cells, key functional sites on the viral genome including cleavage sites for viral proteases PL-PRO and 3CL-PRO as well as cleavage sites for host proteases, locations of important RNA secondary structures, the locations of the transcriptional regulatory sequences, locations of protein phosphorylation and glycosylation sites, identification of sites in the virus that are highly conserved or rapidly evolving in closely related viruses in bats and other mammals, as well as a lively “crowd-sourced annotation” set where any researcher can point out additional sites on the viral genome of special functional, diagnostic, or therapeutic significance [59]. This helps researchers to quickly determine if alternate alleles they believe characterize a new viral clade may be significant beyond their role as epidemiological markers. Instructions for producing custom genome-browser tracks for a given phylogeny and variation dataset are provided in Text S4.

### Phylogenetic Uncertainty and Facilitating Tree Comparisons Across Analyses

In the second part of this work, we address concerns arising from phylogenetic uncertainty. As expected for a relatively slowly-evolving and rapidly expanding viral population [60], there is substantial uncertainty in the SARS-CoV-2 phylogeny. This extends well beyond the typically localized impacts of lab-associated and highly recurrent mutations, and instead derives from the fact that most branches in the SARS-CoV-2 phylogeny are supported by few mutations. Undoubtedly, thousands of unique phylogenies will be produced by groups studying this viral outbreak and these may sometimes support conflicting evolutionary relationships. We therefore sought to provide tools to facilitate interpretation of commonalities and differences among such large phylogenies.

### A tree comparison algorithm using entropy-weighted matching splits

There are many metrics for measuring the total distance between two or more phylogenetic trees [61–66]. One popular metric (Maximum Cluster distance (MCdist)) also identifies the best-matching clades between the two trees. A clade in a rooted tree splits the leaf nodes of that tree into two sets: those inside the clade and those outside the clade. Given a clade C in tree T and a clade C’ in tree T’, the split distance between C and C’ is the number of leaves that have to be moved so that the split for C in T becomes equal to the split for C’ in T’. The (nonsymmetric) correspondence between the clades of T and the clades of T’ established by minimizing the split distance is referred to as the “maximum cluster alignment” or “best split alignment” from T to T’, [61]. This is particularly appealing here because we aim to facilitate comparisons across phylogenetic trees both globally and at individual clades.

We implemented a modified version of MCdist in https://github.com/yatisht/strain_phylogenetics to compare two trees, T and T’, both restricted to the same set of samples, with two improvements. First, we proportionally weighted the split distance between each clade C of T to the best matching clade in T’ by the entropy of C, *i.e.,* by H(p) = -p log^2^(p) - (1-p) log^2^(1-p) where p is the fraction of leaves from T that are in C (see Methods, Figure 9). The entropy-weighted matching split distance emphasizes the importance of the clades in T in terms of how much information about the leaves they carry, which helps highlight clades where the most dramatic changes have occured. The sum, over all clades in T, of the entropy-weighted matching split distance to the best-matching clade in T’ is referred to as entropy-weighted total distance from T to T’. Second, we label all internal branches in T and T’, and identify the most similar branches in both trees based on the clades they define. When multiple branches in T’ match the branch b in T with the same best split distance, we report all best-matching branches (Methods). Additionally, we confirmed that our statistic is a robust measure of tree distance, judging by the strong correlation with other frequently used tree distance metrics (Text S3). Our implementation can compute this statistic for two trees of size 10,000 leaves in just 20 minutes on a single CPU, so it scales to the large trees required for SARS-CoV-2 phylogenetics.

**Figure 9:**
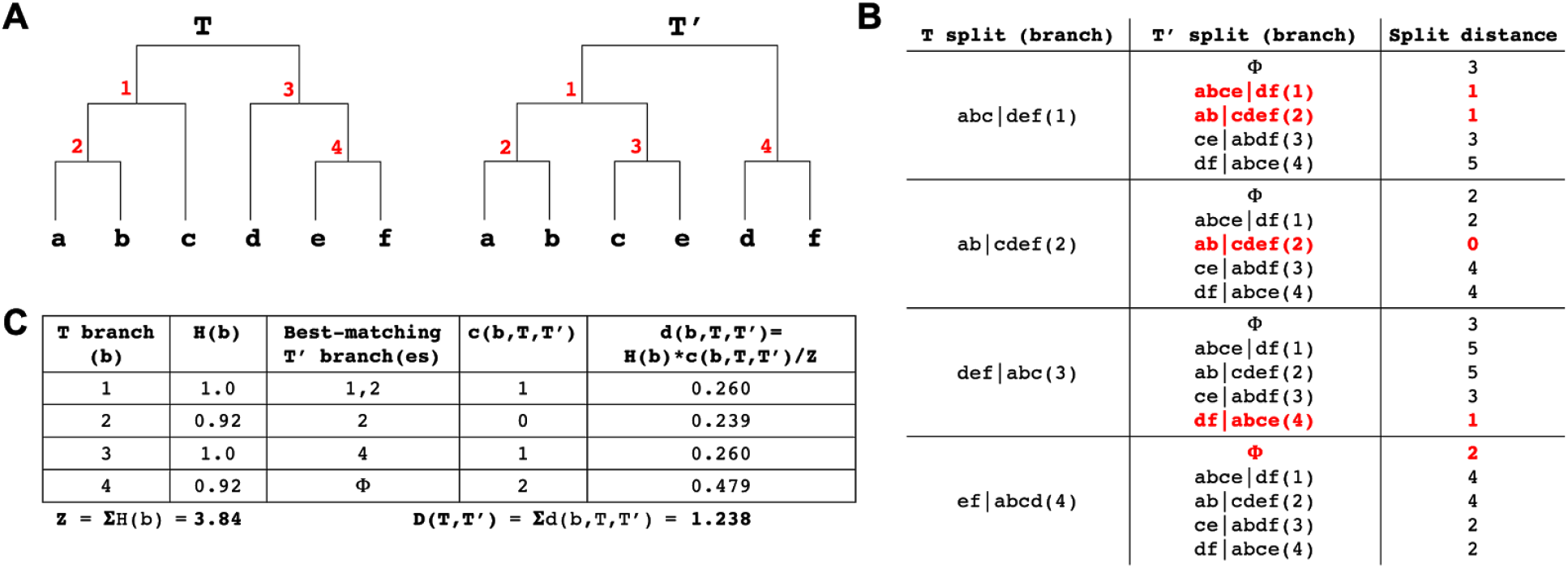
Entropy-weighted distance statistic. **(A)** Example trees (T and T’) for this comparison with identical sets of leaves but different topologies. Internal branches are labelled in red. **(B)** The split distance statistic for each T node (see Methods for notation). Split distance of each T split (branch) from all T’ splits plus a “garbage node” (ɸ) containing a null set of leaves, with the matching split distance and its corresponding T’ split (branch) for each T split (branch) highlighted in red. Multiple T’ splits can match a T split but the garbage node is given precedence (as is the case in T branch 4). **(C)** Table showing the entropy, best-matching T’ branch(es), matching split distance and entropy-weighted matching split distance for each branch in T, as well as the entropy-weighted total distance D(T,T’) between T and T’.

### A Fast Algorithm for Producing Tanglegrams for Trees with Thousands of Leaves

A tanglegram is the most often used method of visualizing the topological difference between two rooted phylogenetic trees defined on the same set of leaf taxa (here, termed samples)[67]. We expect that tanglegrams will have a wide use for analyzing and comparing different SARS-CoV-2 phylogenies. Tanglegrams plot two trees side-by-side with their common leaves connected by straight lines (*e.g.,* Figure S5). For visually appealing and informative tanglegrams, clades in both trees are arranged in a similar vertical order (given the tree topology constraints) with minimum crossing of connecting lines with each other. While there are a number of tree node “rotation” algorithms that optimize tanglegrams for visual appeal [67,68], we found none of the available implementations that we tested [68,69] worked reasonably for phylogenies as large as SARS-CoV-2 phylogenies, either producing unacceptable results or not able to finish the computation. We therefore developed a fast heuristic approach that produces vastly improved tanglegrams (Methods, Figure S5, https://github.com/yatisht/strain_phylogenetics). Our approach takes approximately one minute for the tanglegrams we show here, and we use this heuristic for displaying tanglegrams throughout the text.

### Nextstrain Phylogenies Vary Significantly Over Time

We next explored differences among trees made by the same group from slightly different sample sets with the goal of understanding phylogenetic stability as new samples are incorporated. For the purposes of comparison, we restricted 31 Nextstrain trees produced between March 23, 2020 and April 30, 2020 to just the 468 samples they all have in common. Comparing topologies, we found that a number of these 468 samples moved back and forth between different clade designations during the month (Figure S5), including samples in the specific clades (A1a, A2, A2a, A6, A7, B, B1, B2, B4) named and analyzed by the Nexstrain consortium during this period (*e.g.*, Table S6). Note that the Nextstrain clade ID system was updated while we were finalizing this work [70]. We then measured all pairwise tree distances between restricted trees and found that they varied widely (normalized entropy-weighted total distances ranged from 0.089 to 0.352, Figure 10). There is therefore substantial variation in Nextstrain phylogenies over time.

**Figure 10.**
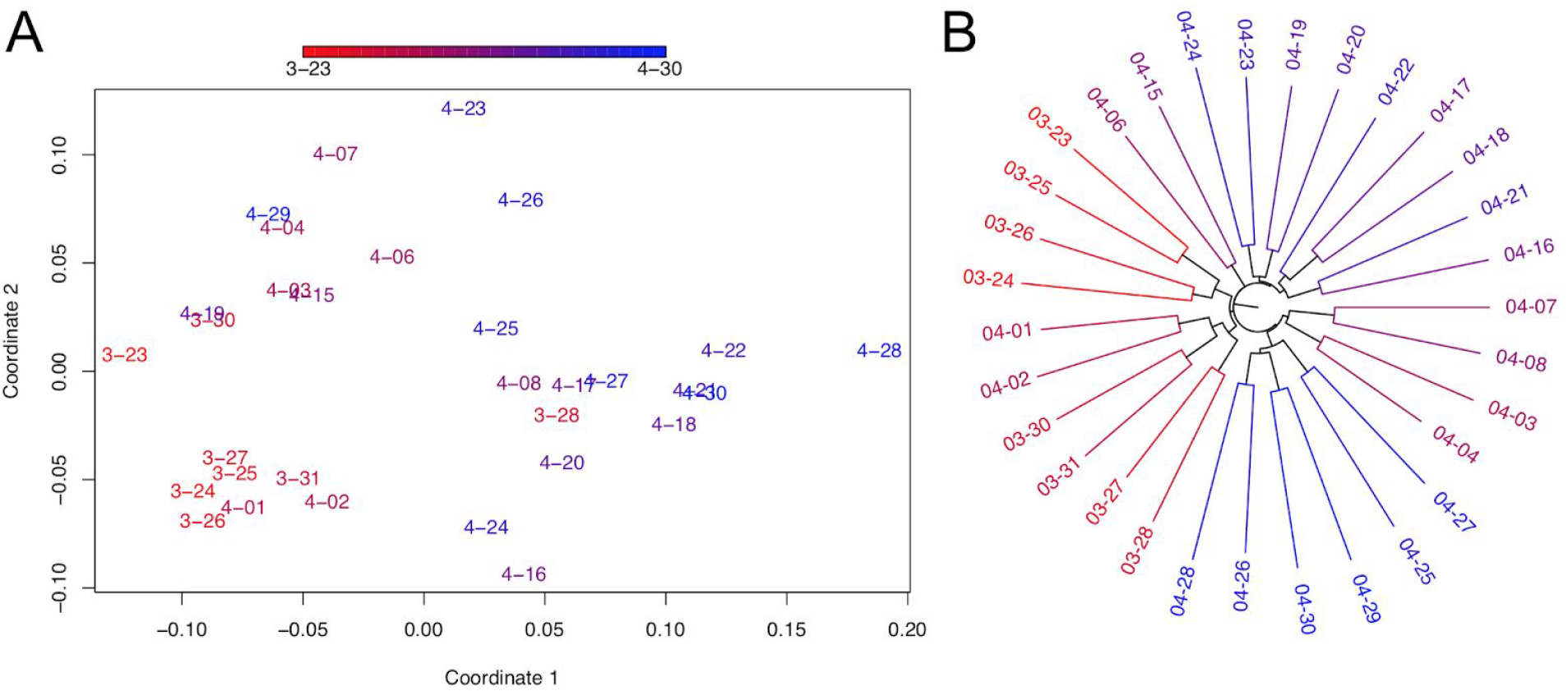
Comparisons of Nextstrain trees over time. (A) Multidimensional scaling of normalized entropy-weighted total distances among phylogenetic trees produced by Nextstrain from March and April. Each topology is labelled with its date and dates are depicted in a color gradient from 3/23 (red) to 4/30 (blue). Coordinates 1 and 2 are plotted here and each contributes 34% and 15% of the total variance explained, respectively. (B) Relationships between Nextstrain phylogenies are shown in a tree-of-trees, “meta-tree” [64] we constructed, which displays the distances among topologies of the constitutive trees. .

Multidimensional scaling (MDS) of the pairwise distances among each topology, as well as meta-tree analysis [64] reveals a strong relationship between topologies and the date that each tree was produced (Figure 10). In particular, the first MDS coordinate is strongly correlated with the release date of the tree (Spearman’s rho = 0.688, P = 3.087e-05). This effect is expected and likely driven, at least in part, by the impact of the sample set used to produce the resulting tree, which necessarily changes as new data are incorporated. Indeed, the proportion of overlapping samples used in constructing each pair of trees is strongly negatively correlated with the normalized entropy-weighted total distance between their topologies (r = −0.384, P = 4e-05, Mantel test), while the set of 468 samples for which we analyze topology is held fixed for all trees. These tools provide the research community a method for tracking the phylogenies of SARS-CoV-2 as the pandemic progresses and phylogenies are produced for larger and larger sample sets. The tools can detect when older clades are confirmed as new samples accumulate, stabilizing inference of these clades, as well as track new subclades as they grow. If inconsistent data is causing persistent clade instability, which may result from lab-associated sequencing errors or actual recombination, it should be visible in this analysis.

### Higher-Level Branches Are Remarkably Consistent Across Analyses

Even if it was possible to obtain error-free data and multiple alignments as well as have all groups use that same data, different tree inference approaches can produce different topologies. Furthermore, there is substantial uncertainty inherent to SARS-CoV-2 evolution because there are few mutations that uniquely mark each branch. Nonetheless, it is essential that epidemiologists studying the pandemic be able to communicate phylogenetically informed observations [17,42]. As discussed above, the clade placements of individual samples, even when inferred by the same group, can vary as different datasets are incorporated into the tree construction process (*e.g.* Table S4, Figure 10). Differences between groups are expected to be even more pronounced. This threatens to leave the community with a “tower of Babel” problem in clade characterization and naming from various different phylogenetic trees. Indeed, the names used for Nextstrain consortium clades (A1a, A2, A2a, A6, A7, B, B1, B2, B4) have nothing whatsoever to do with the clades names (A, B, A.1, A.2, B.1, B.2, A.1.1, etc.) suggested by the COG-UK consortium [17,18], and without a 1-1 correspondence between the topologically defined clades in their respective phylogenetic trees, it is difficult to translate nomenclature in order to conduct precise scientific discourse pertaining to the evolutionary conclusions reached by these groups. Adding further difficulty to this situation, clade naming approaches based on phylogenies must themselves be subject to change as the pandemic spreads and as the evolution of new genotypes requires naming new clades and modifying existing clades. As clade based comparisons are an essential part of consistent scientific discourse, tools are needed to ameliorate these difficulties.

To explore the differences among available phylogenies and to provide guidelines for clade-based comparisons across possible evolutionary histories, we used our approach to identify the correspondence between the Nextstrain phylogeny produced on April 19, 2020 and the COG-UK phylogeny produced on April 24, 2020 (Figure 11A). We observe good agreement between the big Nextstrain named clades and their corresponding best matching named clades in the COG-UK tree and vice versa (*e.g.,* “A2a” clade in Nextstrain, “B.1” clade in COG-UK, etc, Figure 11B), suggesting that these clades are reasonably stable across different analyses. However, in small named subclades within those big clades, there are many noteworthy differences between the two topologies, and the overall congruence is significantly reduced (Figure 11A). In addition to differences in methodology, this reflects a difference in the time when clades were originally named and the intents of each nomenclature system. Nextstrain named clades much earlier and many did not increase in size subsequently, others have since emerged and were named by COG-UK later. Additionally, the COG-UK system is intentionally dynamic and clades that have become inactive are removed. As a consequence, some clades do not have an obvious named analog in the two systems resulting in low similarities (Figure 11B).

**Figure 11:**
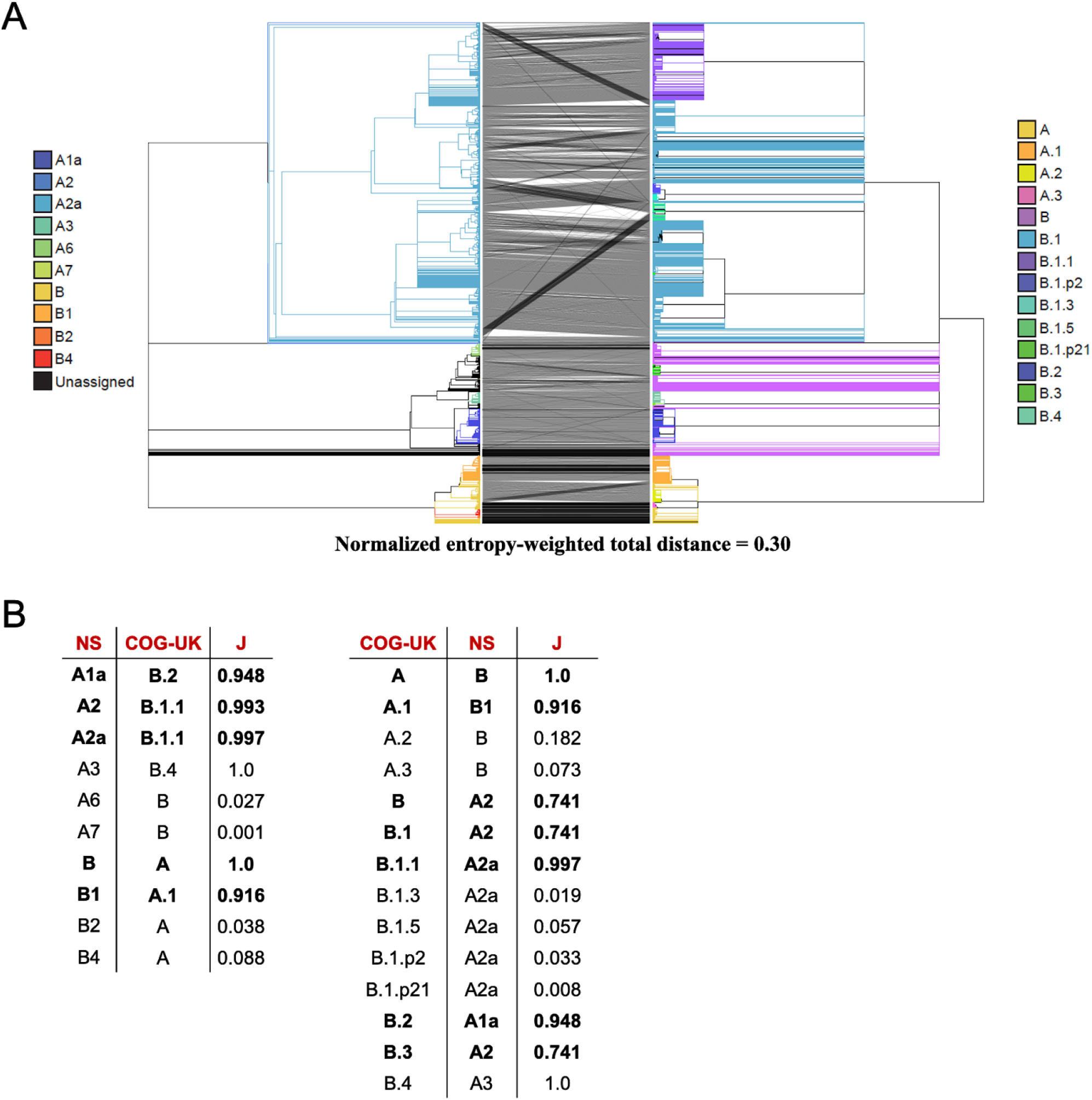
Comparison of Nextstrain and COG-UK trees. **(A)** A tanglegram of the Nextstrain tree from 4/19 (left) with the COG-UK tree from 4/24 (right). Each tree has 4167 samples. **(B)** The COG-UK clades (which they term “lineages”) having the highest Jaccard similarity coefficient (J) with each Nextstrain (NS) named clade and vice versa, where the Jaccard similarity coefficient is computed using the set of samples from the root of that clade. Clades with more than 200 samples are shown in bold font and called “big”, the others “small”. While the naming schemes differ, for each big Nexstrain clade there is a closely corresponding COG-UK clade, and vice-versa.

Perhaps the most obvious difference between the topologies is that the COG-UK tree has many more large polytomies (Figure 11A). This reflects the decisions motivating their analysis [17,71], where the authors’ goal is to provide a well-supported and stable topology to facilitate lucid communication about viral lineages for evolutionary as well as epidemiological studies. This contrasts with the Nextstrain consortium’s primary goal of up-to-date transmission tracing. As is typical in phylogenetics, topological stability comes as a tradeoff against the cost of articulation in the branches. Because of the many different motivations for constructing phylogenetic trees, it is a certainty that many independent trees will be used to study the evolution of SARS-CoV-2. Comparisons using our approaches can enable communication about evolving viral lineages across disparate analyses by facilitating the identification and visualization of the most closely matching clades.

### Higher Branches in Our Tree Closely Mirror A Nextstrain “Consensus” Tree and the COG-UK Tree

To identify stable nodes across analyses we compared a Nextstrain “consensus tree” and the COG-UK tree. To do this, we produced a majority rule clade consensus tree [72] for the 422 common samples in 31 Nextstrain releases between 3/23 to 4/30, and restricted the COG-UK tree to these same samples. We find exceptionally good congruence between our Nextstrain consensus and the COG-UK phylogenies (Figure 12A), even though the inference methods differed substantially. Specifically, the COG-UK tree is built using a more typical bootstrapping approach [58] whereas our approach for building a Nextstrain “consensus” from trees produced on subsequent days would resemble a kind of “bootstrapping by samples” approach. This congruence reaffirms the idea that the COG-UK tree provides a stable “backbone” to enable direct conversations in epidemiology. Nonetheless, we still observe several small rearrangements between the two topologies, suggesting that both will likely be subject to clade refinements in the future.

**Figure 12:**
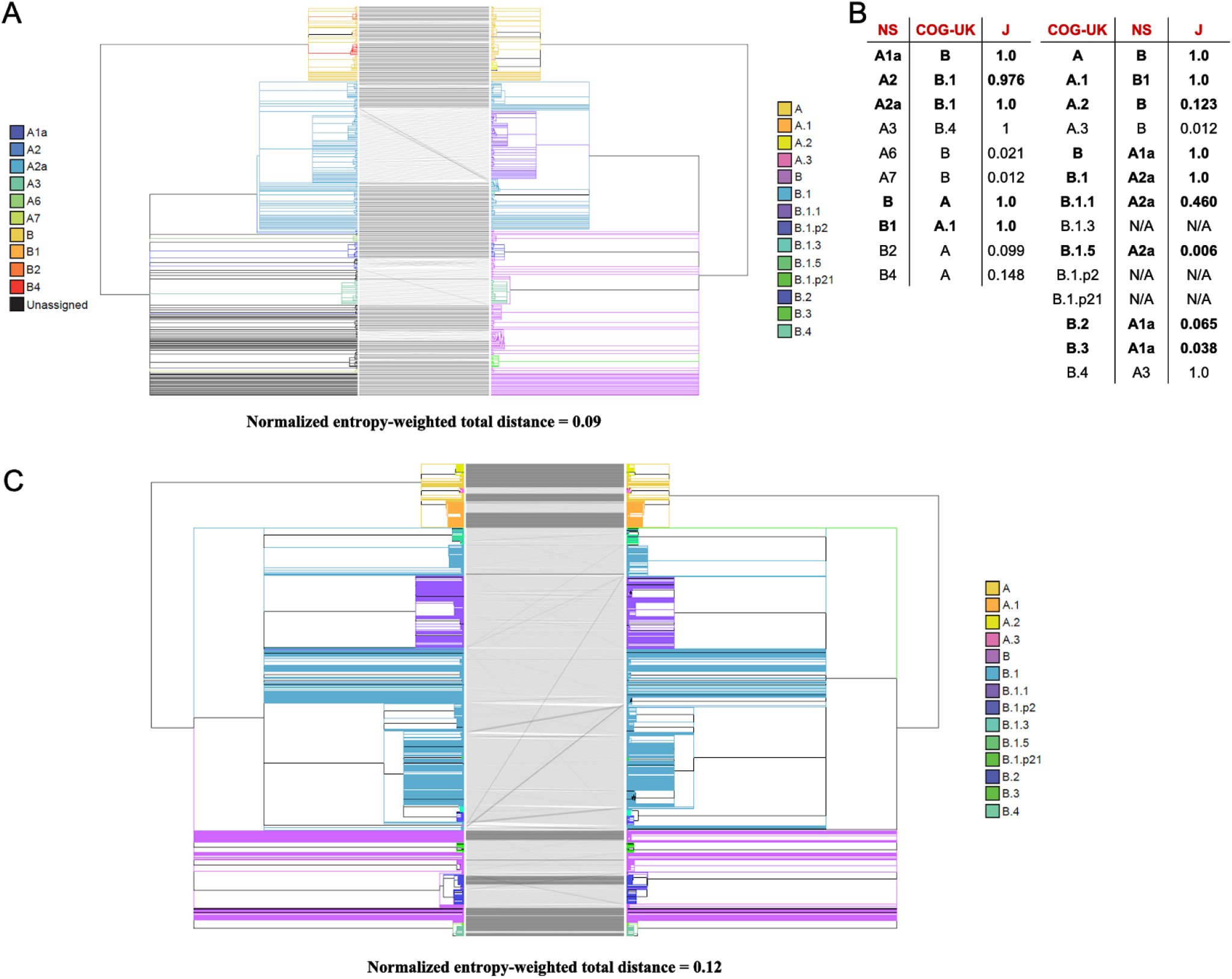
Comparison of Nextstrain and the COG-UK trees. **(A)** A tanglegram of our Nextstrain consensus tree (left) and COG-UK tree from 4/24 (right). Each tree has 422 samples. **(B)** The COG-UK lineages having the highest Jaccard similarity coefficient (J) with each Nextstrain consensus (NS) named clade and vice versa. Big clades defined in Fig. 11 (those containing 200 or more samples in the Fig. 11A trees) are in bold. Lineages in ‘N/A’ (B.1.3, B.1.p2 and B.1.p21) were pruned out as a result of restricting the trees to common samples. **(C)** A tanglegram of our tree produced after masking all lab-associated and extremal mutations except 11083 (left) and COG-UK tree from 4/24 (right). Each tree has 4172 samples and the samples (branches) have been colored based on COG-UK lineage labels.

We also observed good overall congruence between the tree that we produced after removing lab-associated and extremal mutations (except 11083, see above) and the COG-UK tree (Figure 12C). Here, the sample size is much larger, 4172, allowing for a much more quantitative comparison. The correspondence between the two trees is very high with normalized entropy weighted total distance of just 0.12. Because lab-associated and extremal mutations were used in the COG-UK tree but not in our tree, this consistency among topologies supports our assertion that the effect of lab-associated and extremal mutations will typically not result in large-scale reorganizations of large clades across the phylogeny. Each tree including our Nextstrain “consensus” is available for visualization through the UCSC Genome Browser (Figure 8, S6).

### Powerful Tools for Visualizing, Interpreting Differences Among Phylogenies

Different analysis goals require varying levels of phylogenetic resolution and certainty, and it is very likely that hundreds of partially independent phylogenies will be produced studying SARS-CoV-2 evolution. For that reason, we have sought to provide the community with effective methods for tree-based comparisons. In particular, here we provide (1) improved methods for quantitative comparison among trees at the level of whole topologies and at individual nodes; (2) an extremely rapid tanglegram clade rotation method for visualization of differences among tree topologies; and (3) dynamic tree visualization capabilities within the SARS-CoV-2 Genome Browser. Importantly, each method that we present scales well to thousands of samples, and is integrated into the SARS-CoV-2 Genome Browser to facilitate rapid comparison with existing phylogenetic datasets, and to cross-reference sites to molecular information relevant to basic biology, diagnostics, and therapy. Software to run each analysis is available from https://github.com/yatisht/strain_phylogenetics.

## Conclusion and Outlook

The SARS-CoV-2 pandemic has driven an impressive global community response providing real-time sequencing data to trace the viral outbreak [1–5]. Because these efforts are both decentralized and urgent, there is potential for systematic differences in data generation and processing to inject inappropriate biases and signal into these data [24]. Similarly, thousands of distinct and differing phylogenies will be made from these data. In this work, we sought to provide tools to detect and interpret sources of conflict and uncertainty in local and global phylogenies. We integrate these into powerful visualization systems to facilitate continued global analysis of viral population dynamics.

## Methods

### Obtaining Nextstrain Trees and Genotype Data

We have downloaded genomic variation data from http://nextstrain.org/ncov, which is ultimately processed and derived from the GISAID database [73], and transformed it into Variant Call Format (VCF, [74]) file with genotypes for all samples as assigned by Nextstrain, a Newick tree file, and associated files for display in the UCSC SARS-CoV-2 Genome Browser. Software to perform this is described here (https://github.com/ucscGenomeBrowser/kent/blob/master/src/hg/utils/otto/nextstrainNcov/nextstrain.py).

### Obtaining and Correcting Sample Metadata

We obtained the GISAID metadata table in bulk from GISAID [75]. Before we were able to search for lab-associated mutations, we identified various errors in GISAID metadata files, most of which appear to be due to misspellings and inconsistent naming conventions of “originating” and “submitting” labs across separate sample submissions. We therefore developed a simple approach to detect these errors systematically based on the character content and length of “originating” and “submitting” lab names (https://github.com/lgozasht/COVID-19-Lab-Specific-Bias-Filter). We merge coincident metadata under consistent lab names if “originating” or “submitting” lab names share 70% length similarity and 90% character similarity or 70% length similarity and 80% identical character positions, and output a revised metadata file. We checked all merged names by hand to ensure accuracy, and we maintain a log of each merger event and annotate low confidence mergers. Our updated metadata table is available from https://github.com/lgozasht/COVID-19-Lab-Specific-Bias-Filter.

### Identification of Highly Recurrent Mutations

To detect mutations that reoccur many times through viral evolution, we computed the parsimony score [45,46] for each polymorphic site (our program is available from https://github.com/yatisht/strain_phylogenetics). Briefly, conditional on a tree, we compute the minimum number of branches that have experienced a mutation at a single site to accommodate the phylogenetic distribution of the mutant and reference allele. These are candidate highly recurrent mutations, but we note that these mutations, or others elsewhere on the chromosome, might also impact the process of tree building itself, and the score should be interpreted with caution if counting the specific rate of occurrence at a given site is of interest.

### Automated Identification of Extremal sites

After computing the parsimony score for each polymorphic site, we identified a set of extremal sites that displayed exceptional parsimony scores relative to their allele frequencies as follows. First, we excluded sites with rare alternate alleles, *i.e.* sites whose alternate allele frequency was found to be lower than a certain threshold K, where K is the maximum alternate allele frequency at which at least two sites had saturated parsimony scores (*i.e.* parsimony score equals alternate allele count). Second, we extracted sites whose parsimony score was found to be the highest among sites with the same or smaller alternate allele frequency. Finally, we also required that extremal sites have an alternate allele frequency that is lowest among all sites with its parsimony score or higher. A program to perform this search is available at https://github.com/yatisht/strain_phylogenetics. This program also optionally allows for extremal sites to be identified without including C>U mutations as these are particularly abundant in SARS-CoV-2 genomes.

### Discovery of Lab-Associated Mutations

We systematically flagged possible variants resulting from lab-specific biases based on the proportion of lab-specific alternate allele calls and respective alternate allele frequency (https://github.com/lgozasht/COVID-19-Lab-Specific-Bias-Filter). To do this, we first filtered variants with parsimony score greater than 4 using concurrent Nextstrain tree and vcf files from 4/19/2020. Next, we obtained metadata for all COVID-19 genomes on GISAID (accessed 4/28/2020) and computed the proportion of alternate allele calls contributed by each “originating lab” and “submitting lab” for each filtered variant. We then employed a Fisher’s exact test associating the number of major and alternate alleles attributed to each specific “originating” and “submitting” lab and the respective global major and alternate allele counts. We flagged variants for which one lab accounts for more than 80% of the total alternate allele calls and for which a Fisher’s Exact Test suggests a strong correlation (at the p < 0.01 level) between that lab and samples containing the alternate allele. We note that these cutoffs are somewhat arbitrary, and may require modification in the future, but the subdivision of the data is consistent with our expectations as described in Results. Because samples are not independent and identically distributed, p-values may not reflect error but rather relatedness among samples sequenced at a single facility. For example, if a single lab sampled a transmission chain, many mutations could be strongly associated with that facility. These should be interpreted cautiously, however, there is no obvious reason why unrelated samples sequenced at the same facility should share an excess of homoplasious mutations.

### Testing for Overlap with ARTIC Primers

To compare our highly recurrent mutations to the ARTIC primer set, we downloaded the positions of the ARTIC primer binding sites from (https://github.com/artic-network/artic-ncov2019/blob/master/primer_schemes/nCoV-2019/V3/nCoV-2019.bed, last accessed 5/6/2020). We computed the number of mutation in each category that overlapped primer binding sites, and we computed the mean distance between each variant and the nearest primer binding site. To test for enrichment for overlap and proximity to primer binding sites, we performed a permutation test where we selected positions at random without replacement across the viral genome to compare to our observed distribution for the real mutations. Each permutation was performed 10,000 times.

### A Clade Comparison Method using Branch Splits

Comparison of clades is made using a symmetric notion of a clade that are called **splits**as defined in TreeCmp [61]. In a rooted tree, the branches are directed to point away from the root, and a directed branch defining a clade divides all the leaves (lineages) into 2 categories: those in the clade (reachable by following additional directed edges forward from the branch; we call this being “inside” the branch) and the rest, *i.e.* those not in the clade (“outside” the branch, we might say these samples are in the “unclade”). It is the root of the tree that polarizes each split by providing a direction for the branch; *i.e.* providing a concept of “inside” versus “outside”, or equivalently “clade” versus “unclade”. These two sets, say A and B, define the split. The split is denoted as A|B.

Two phylogenetic trees are similar if their branches produce a similar set of splits. When comparing two phylogenetic trees, we begin by finding the common leaf set. That is, the set of leaves (lineages) that are included in both trees. Then for each tree and each branch in that tree, the **reduced split**is obtained from the split by removing all samples not in the common leaf set for the two trees being compared. To compare two reduced splits, A|B and X|Y, we first compute the size of the set-theoretic symmetric difference between the clades A and X, *i.e.*, the number of samples that are in A but not in X (denoted by |A\X|), plus the number of samples that are in X but not in A (denoted by |X\A|). This number is denoted by s(A|B,X|Y) and is called the **split distance** between the reduced splits A|B and X|Y. Symbolically

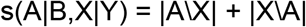

The same comparison of B with Y is not necessary as it will yield the same number as obtained by comparing A and X.

Now, if b is a branch in tree T and A|B is its reduced split, the **matching split distance** of the branch b in tree T’ is

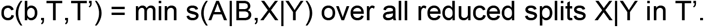

Given the reduced split A|B for a branch b in a tree T and the set of all reduced splits in a second tree T’, *i.e.* {X|Y : X|Y is a reduced split in T’}, the set of **best matching splits** for A|B is in T’ is defined as

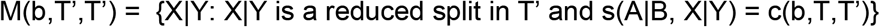

That is, every reduced split in M(b,T,T’) has a split distance from A|B equal to the matching split distance of branch b in T’, which is the smallest distance possible. The branches corresponding to the best matching splits are called **best matching branches**.

We can also define the (Shannon) **entropy** of the branch b in the tree T as the entropy in units of bits of its reduced split A|B. Let p = |A|/(|A|+|B|) where |S| denotes the cardinality of the set S.

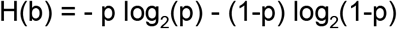

The proportional entropy weight of the branch b in the tree T is the normalized entropy

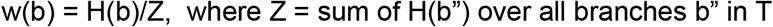

The **entropy-weighted matching split distance** to tree T’ of branch b in tree T is

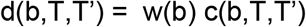

We define a distance measure, called **entropy-weighted total distance**, for two trees T and T’, as the sum of entropy-weighted matching split distance for all branches in T:

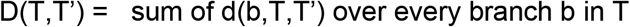

As this distance measure is not symmetric, we also define a **symmetric** version of it as

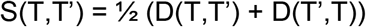

Since the above metric scales with the size of trees being compared, we also define a **normalized** version using the expected distance [76], which is computed using trees T^p^ and T^p^’ that randomly permute the leaves of T and T’, respectively, while maintaining the tree structure, as

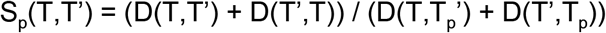

Code for computing these distance measures can be found at https://github.com/yatisht/strain_phylogenetics. This code has additional features, such as the ability to replace the Shannon entropy - p log^2^(p) - (1-p) log^2^(1-p) with related weighting functions such as 2 min{p,1-p}. We find that the method is robust to such replacements (data not shown).

### Clade Orientation for Tree Comparison

While node rotation algorithms in the context of tanglegram visualization have been implemented in the cophylo and Dendroscope3 tools [67,68], we found these algorithms to be either too slow or inadequate for the large SARS-CoV-2 phylogenies that we compared. We implemented a simple node rotation heuristic, RotTrees, that works well and completes in reasonable time (~1 min) for SARS-CoV-2 trees with ~5K leaves. The algorithm RotTrees accepts two trees, T and T’, each pruned to only contain the shared set of leaves, as input. First, while maintaining the leaf order of T, RotTrees makes a breadth-first traversal in T’, rotating the children of each traversed node based on its average rank (*i.e.* child with a lower average rank appears higher), which is the average of the positions of the appearance of that child node's leaves in T. Second, RotTrees repeats the above to rotate the leaves of T while maintaining the leaf order of T’. The previous two steps are repeated until convergence (no new rotations in that iteration) and the final tree rotations for T and T’ are returned. We made this routine available in https://github.com/yatisht/strain_phylogenetics. This may not be optimal for all tree co-visualization purposes, but here we find that this approach is sufficient to produce vastly improved tree visualizations than many available packages.

### Phylogenetic Trees

We obtained the phylogenetic tree hosted by Nextstrain (accessed 4/19/2020) and used this in our comparisons of clades among trees and as our primary data object for examining apparently recurrent mutation on the tree. We did separately confirm that most apparently recurrent mutations are recovered on the trees produced on different days by Nextstrain.

For comparison of clades among different tree-building approaches, we obtained variant datasets, and phylogenies from Nextstrain (https://nextstrain.org/ncov accessed 4/19/2020-4/26/2020), and from COG-UK (https://cog-uk.s3.climb.ac.uk/20200424/cog_2020-04-24_tree.newick, accessed 4/24/2020)

### Phylogenetic Reconstruction

From the 04/19 Nextstrain release, we created a “reference phylogeny” using IQ-TREE-2 [57,77] to build phylogenies from each of these alignments using the GTR+G nucleotide substitution model. For all other phylogenies, we altered the input by removing or “masking” individual sites, then produced phylogenies from these altered alignments using the same IQ-TREE-2 parameters.

The likelihood of a tree given the alignment from which it was constructed was automatically calculated by the IQ-TREE command used above (*iqtree -s <alignment.phy> -m GTR+G*). However, to compute the likelihood of a particular alignment given a different tree, we used the command *iqtree -s <alignment.phy> -te <phylogeny.nh> -m GTR+G*.

To generate our final tree having masked lab-associated and extremal mutations, we used the same command but also included the ultrafast bootstrapping option “-bb 1000” to assist with quantifying uncertainty in our final phylogeny [58]. We used the same command but included the full multiple alignment to compare the tree obtained to one obtained from the full dataset using identical methodology. Finally we collapsed all branches that were not supported by at least one mutation using parsimony to identify nodes that experienced a mutation.

### Systematic Error Addition Experiments

To investigate the effects of lab-specific alleles on phylogenetic topology, we also introduced artificial errors at control sites. We chose three sites at which to introduce these errors: A11991G, C22214G, and C10029U. To introduce an error, we manually changed a reference allele to an alternate allele for a given sample at a given site. For each of these sites, we chose 10, 25, and 50 samples for which we introduced errors. To mimic the effects of a lab-specific allele, we ensured that each set of samples with artificial errors must come from the same country. We chose Australia due to its high representation in the Nextstrain data, as 372 samples in the 04/19 Nextstrain release came from Australia. To further mimic lab-specific behavior, we separately introduced errors at the same sites for 10, 25, and 50 randomly selected French samples collected between March 1 and March 17. After introducing these errors, we constructed phylogenies from the modified alignments using IQ-TREE 2 [57,77], as described above. In total, we produced 54 phylogenies in this experiment, introducing errors at three sets of random samples for each of the three sites, at 10, 25, and 50 samples each, for Australian and French samples.

We also repeated this experiment, but introducing errors at pairs of sites simultaneously rather than at individual sites (i.e. A11991G and C22214G, A11991G and C10029U, and C10029U and C22214G). We used the same randomly chosen sets of French samples for this aspect of the experiment, and produced phylogenies by the same methods. In total, we produced 27 phylogenies in this experiment, introducing errors at three sets of randomly chosen samples, at 10, 25 and 50 samples each, for each of the three pairs of sites.

### Comparisons Across Nextstrain Trees

To understand commonalities in tree structure over time, we used multidimensional scaling of a distance matrix of normalized entropy-weighted total distances among Nextstrain releases (pruned to 468 shared samples) spanning from March 23 to April 30. To do this, we used the cmdscale() function in base R (https://www.R-project.org/), and we retained the first six coordinates because they accounted for the vast majority of the total variance explained (approximately 80%). We computed the correlation between our distance matrix and the proportion of samples shared among topologies produced each day using a Mantel test implemented within the ade4 package in R.

### Producing a Nextstrain Consensus Tree

To produce a Nextstrain consensus tree we first pruned all Nextstrain trees to a common set of samples included in each tree. We then used the sumtrees script within the dendropy package [69] to produce a majority rule consensus tree out of each tree requiring at least 50% of trees support a clade for inclusion in the final consensus tree. Specifically, we used the sumtrees function to perform this task. In our cases, that is equivalent to requiring at least 16 of 31 trees contain a given clade to include it.

## Acknowledgements

The authors thank the GISAID database, Nextstrain consortium, COG-UK consortium and all labs who contributed SARS-CoV-2 sequence data. We additionally thank Nick Loman and Jared Simpson for feedback early on when we began to notice anomalous data. Additionally, several groups have been extremely forthcoming with information about the likely sources of lab-associated mutations and eager to correct them.

---

**Text S1. High Allele Frequency Variants Could Reveal Cross-Contamination**

Although much of this work is focused on detecting and characterizing the impacts of low-frequency highly recurrent and lab-associated alleles, cross-contamination among samples is also a potential source of widespread phylogenetically discordant sites that warrants mentioning. The majority of labs performing viral genome sequencing are processing multiple samples. There is therefore a significant possibility for contamination to drive the apparent recurrence of high frequency mutations.

Unfortunately, contamination, short recombination tracts, and high frequency recurrent mutations create largely similar predictions about the distributions of recurrent alleles (Table 1). Yet there are some distinctions one might expect. Both recombination and contamination require a sample or lineage to encounter another of a different allele to be observable. Therefore, high frequency alleles should typically be involved in both recombination and contamination events. Additionally, all else being equal, we expect that we would observe equal numbers of forward and backward mutations across the tree for recombination and contamination, but not for recurrent mutation which can be quite biased (see above). Consistent with this idea, six out of eleven sites with alternate allele frequency above 10% show evidence of additional forward and backward mutation even after removing lab-associated mutations (Figure 2, Table S3). Hence, even if we could remove all systematic errors, contamination should be considered as a possible source of homoplasious mutation before confident conclusions are drawn about natural selection or the presence of viral recombination.

**Text S2. Potential for Correlated Error in Our Dataset**

To investigate the potential for highly correlated lab-associated mutations in the real data, we extracted the set of mutations with alternate allele counts of 10 or more and where at least 80% of samples containing alternate alleles were derived from a single lab (Table S4). This set of alleles shares many features with sites that we believe to be sites of real variation, suggesting that many are indeed true variants. In aggregate, these mutations are not enriched for proximity to ARTIC primer binding sites (P = 0.9502, permutation test), the C>U mutation fraction is similar to that observed in high frequency sites (P = 0.5307, Fisher’s exact test), they affect amino acids at similar rates to those of high frequency alleles (P = 0.7643), and they have low parsimony scores even relative to our “two-error” experiments (1-2, Table S4). Our results therefore suggest that highly correlated lab-associated mutations are relatively rare.

It is noteworthy that there is overlap with some of the lab groups who contributed high parsimony lab-associated alleles (nine out of 24, Table S1). However, these are also groups who contributed the most genome sequences in our dataset and they are therefore the most likely to be associated with low frequency variation or lab-associated mutations for that matter. Most such mutations do not overlap in samples with other lab-associated mutation sites indicating that if they were independent mutations we would likely see their placement vary across the tree. Nonetheless, in one extreme case, G11417U, U14073C, and A23947G co-occur in a single clade across many samples suggesting that these sites could impact tree-building algorithms even more substantially than those in our two-error simulations.

However, samples containing these mutations are not unusually divergent relative to the clade size (the average pairwise nucleotide diversity is 1.18 sites/genome) as would be expected if they were incorrectly grouped. Moreover, we emphasize that real variants should be correlated on the viral phylogeny and that these do not constitute sequencing errors. We believe that our approach has likely identified the majority of lab-associated recurrent mutations in this dataset that occur in more than a handful of samples.

**Text S3. Entropy Weighted Distance is a Robust Tree-Distance Measure**

To confirm that our distance measure will be robust and consistent with expectations, we compared the set of all pairwise distances between trees produced by Nextstrain from March 23 to April 30 across a range of tree distance statistics. In particular, we find that entropy-weighted total distance is strongly correlated with quartet, Robinson-Foulds and matching-split tree distance measures (P < 1e-5, in all cases, Mantel test). This strongly suggests that our approach yields robust and interpretable tree distances.

**Text S4. Step-by-step instructions for setting up a genome browser session with a custom tree and VCF**

We provide researchers the ability to upload their own set of aligned SARS-CoV-2 genomes and an accompanying phylogenetic tree. This allows the researcher to compare their tree to trees from Nextstrain and COG-UK, and to map alleles characteristic of particular clades to sites in the virus genome of functional, diagnostic or therapeutic significance. Alignment files built by most tools are too large to transfer when the number of viral samples gets large, so we require they be converted to the more compact VCF format with sample genotypes before upload (the <u>Msa2Vcf</u> tool produces VCF with sample genotypes)‥ Once the VCF file for the alignment is created, one does the following:

1. Compress the VCF file with bgzip and index it with tabix following the instructions here: https://genome.ucsc.edu/goldenPath/help/vcf.html
2. Place the .vcf.gz, .vcf.gz.tbi and a newick format file for the phylogenetic tree (or trees) on a web or ftp server accessible to genome.ucsc.edu. As a hypothetical example, all relevant files could be available from the same server as shown:

https://my.lab.org/my.vcf.gz
https://my.lab.org/my.vcf.gz.tbi
https://my.lab.org/my.newick
3. Replace the example URLs with actual URLs in the following custom track specification line (all one line, no line breaks), copy and paste into the input in https://genome.ucsc.edu/cgi-bin/hgCustom (making sure the SARS-CoV-2 genome is selected):

~~~
track name=myTreeAndVcf type=vcfTabix visibility=pack
hapClusterEnabled=on hapClusterHeight=500
bigDataUrl=https://my.lab.org/my.vcf.gz hapClusterMethod=“treeFile https://my.lab.org/my.newick”
~~~

**Figure S1.**
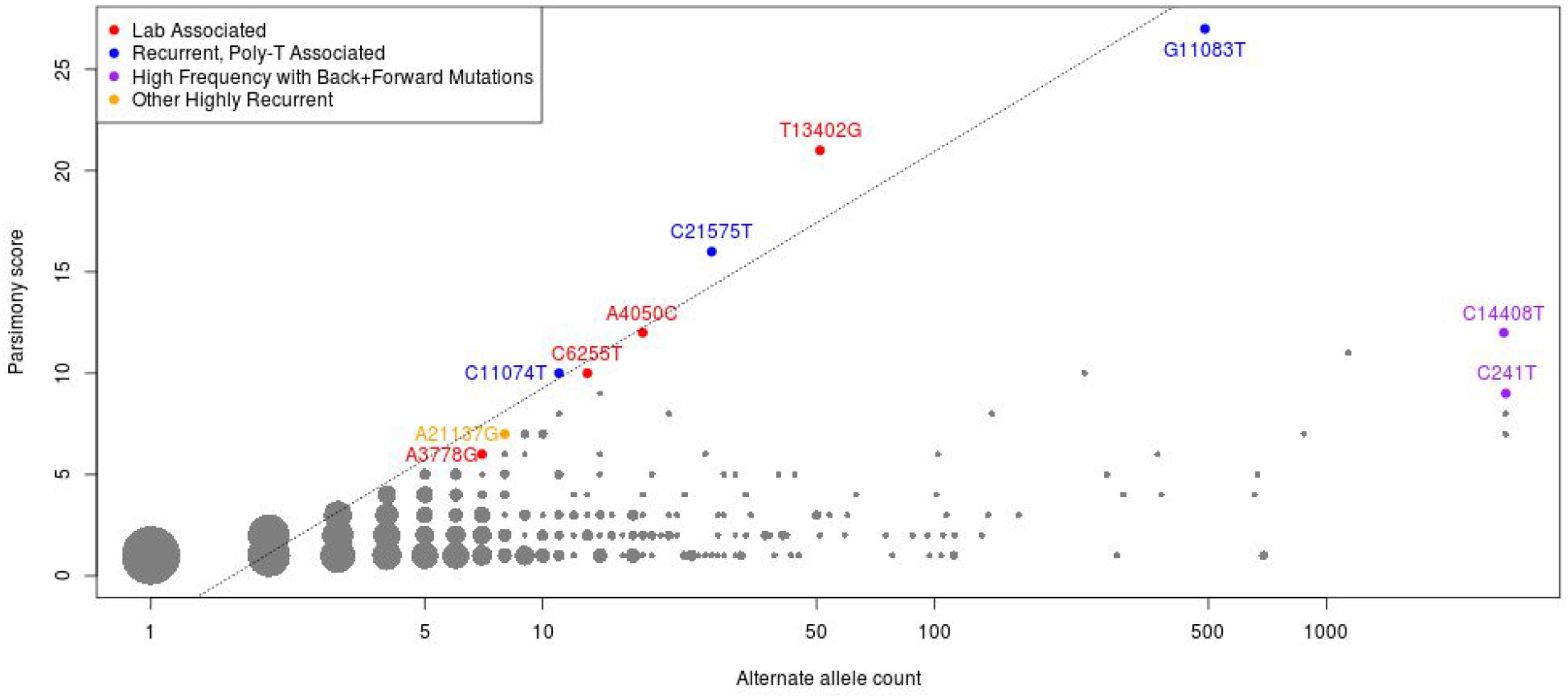
Alternate allele count versus parsimony score for the Nextstrain 4/20/2020 dataset and tree. Each point is labeled as in Figure 2A with additional extremal points annotated. The dashed line is fit to the extremal points and has log2-base slope 3.518.

**Figure S2:**
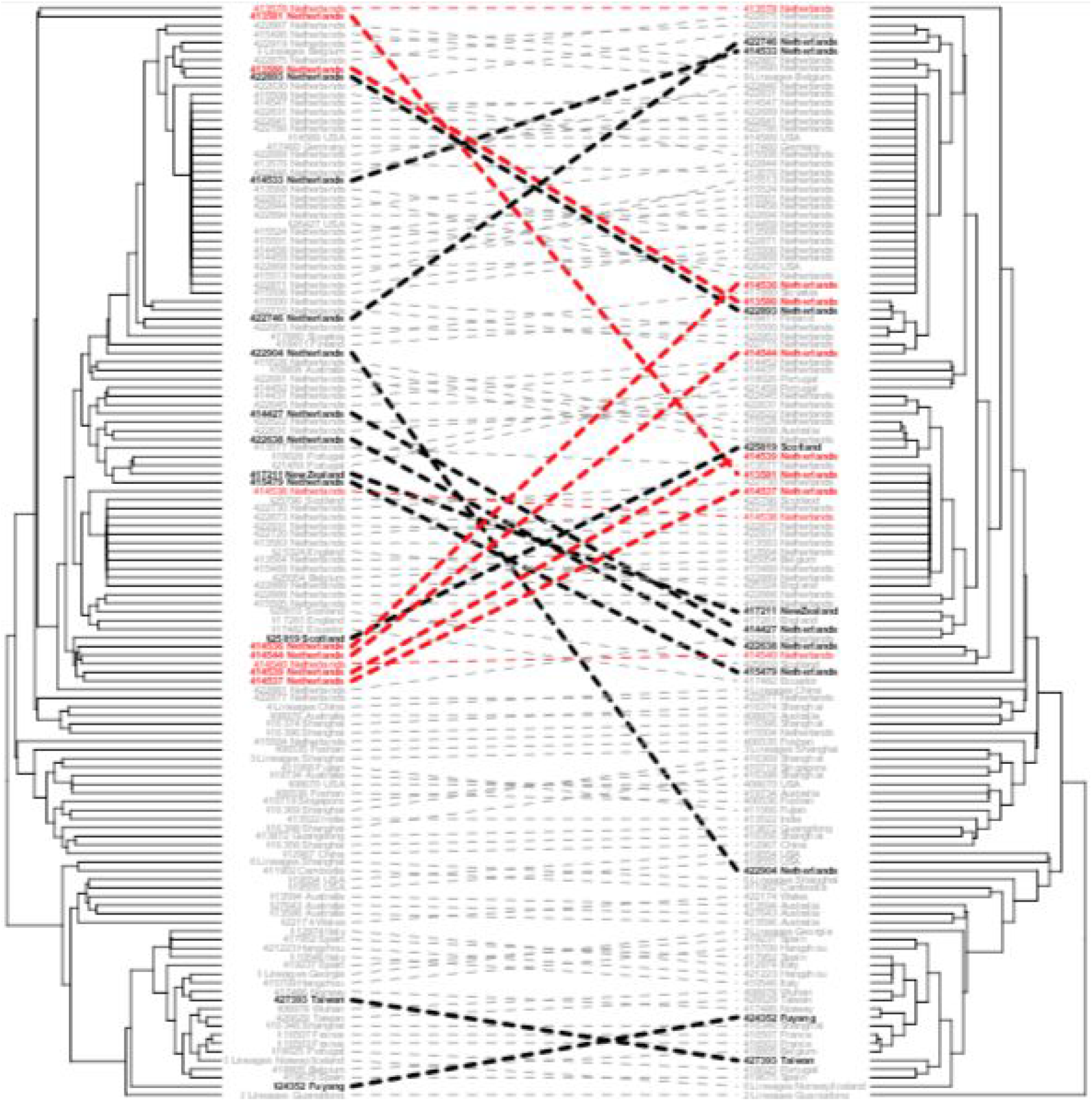
Lab-associated mutations influence tree topology. Phylogenies created using the variants from 04/19 Nextstrain release without modification (left) and with lab-associated mutations completely masked (right) demonstrate movement of multiple samples between sub-clades. Those samples with the greatest changes in placement between the phylogenies are bolded. This includes many samples containing lab-associated mutations that we masked, which are colored in red.

**Figure S3:**
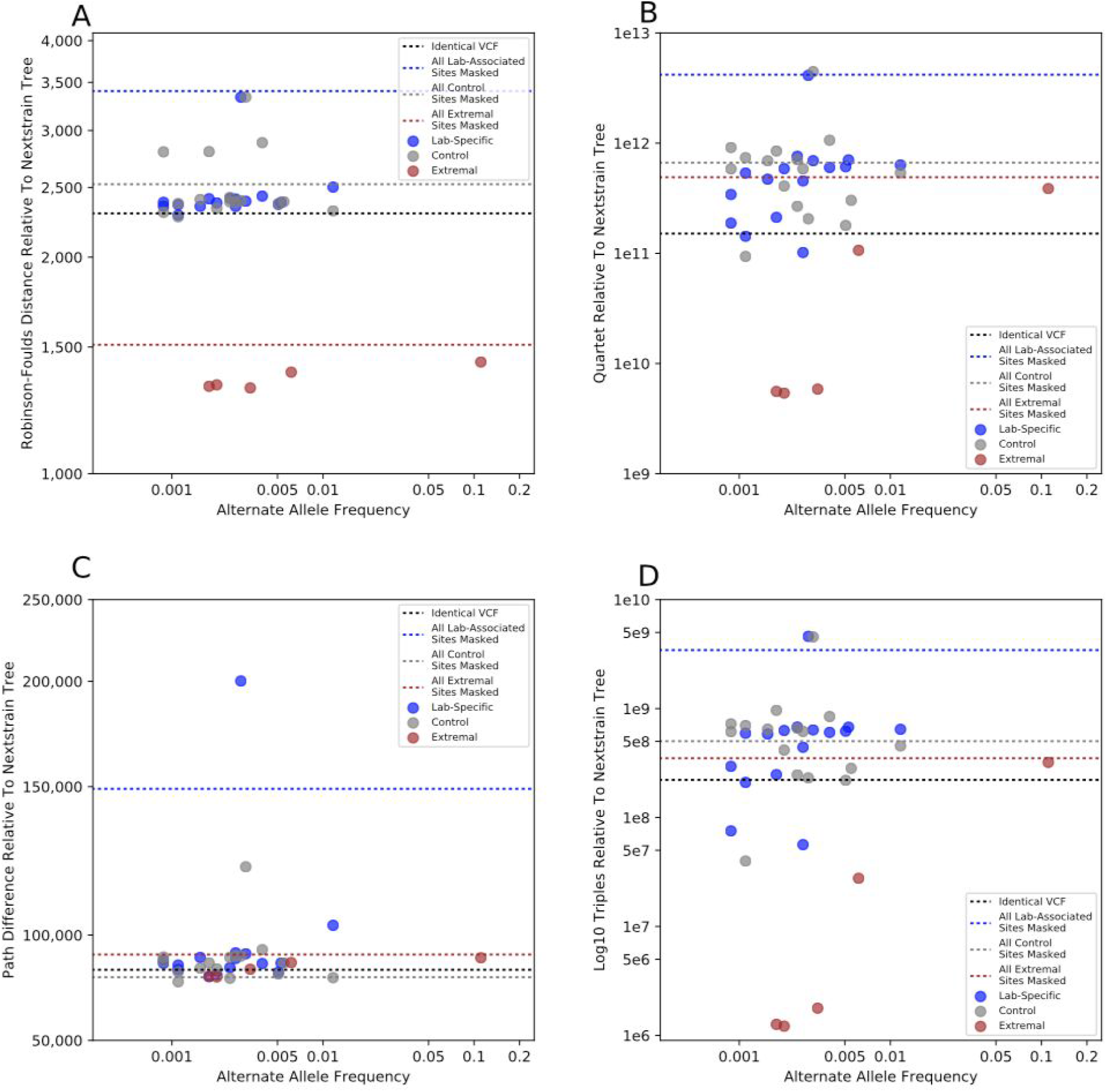
Comparisons between the reference phylogeny, built from the 04/19/2020 release of Nextstrain, to phylogenies built by entirely masking lab-associated mutations (blue), control sites (grey), and extremal sites (brown) are shown for Robinson-Foulds (A), Quartet (B), Path Difference (C), and Triples (D) scores as calculated by TreeCmp [61]. Horizontal lines indicate scores for phylogenies constructed after masking all lab-associated sites (blue), all control sites (grey), all extremal sites (brown), or using an unaltered Nextstrain 04/19/2020 dataset (black).

**Figure S4:**
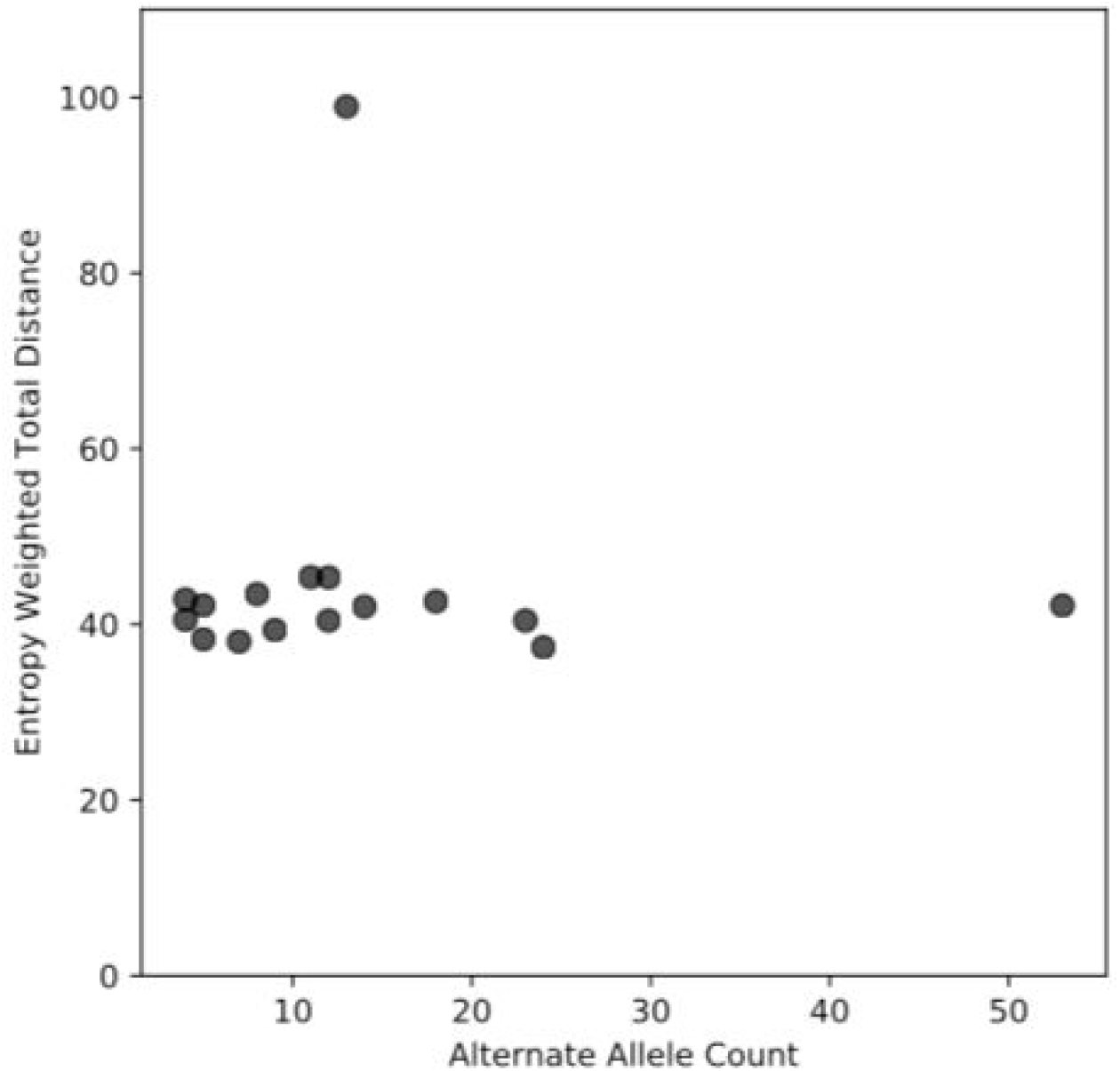
The entropy-weighted total distance values between the reference phylogeny and phylogenies constructed after entirely masking all samples with an alternate allele at a given site are shown. The sites used here are the same sites corresponding to lab-specific shown in Figure 5.

**Figure S5:**
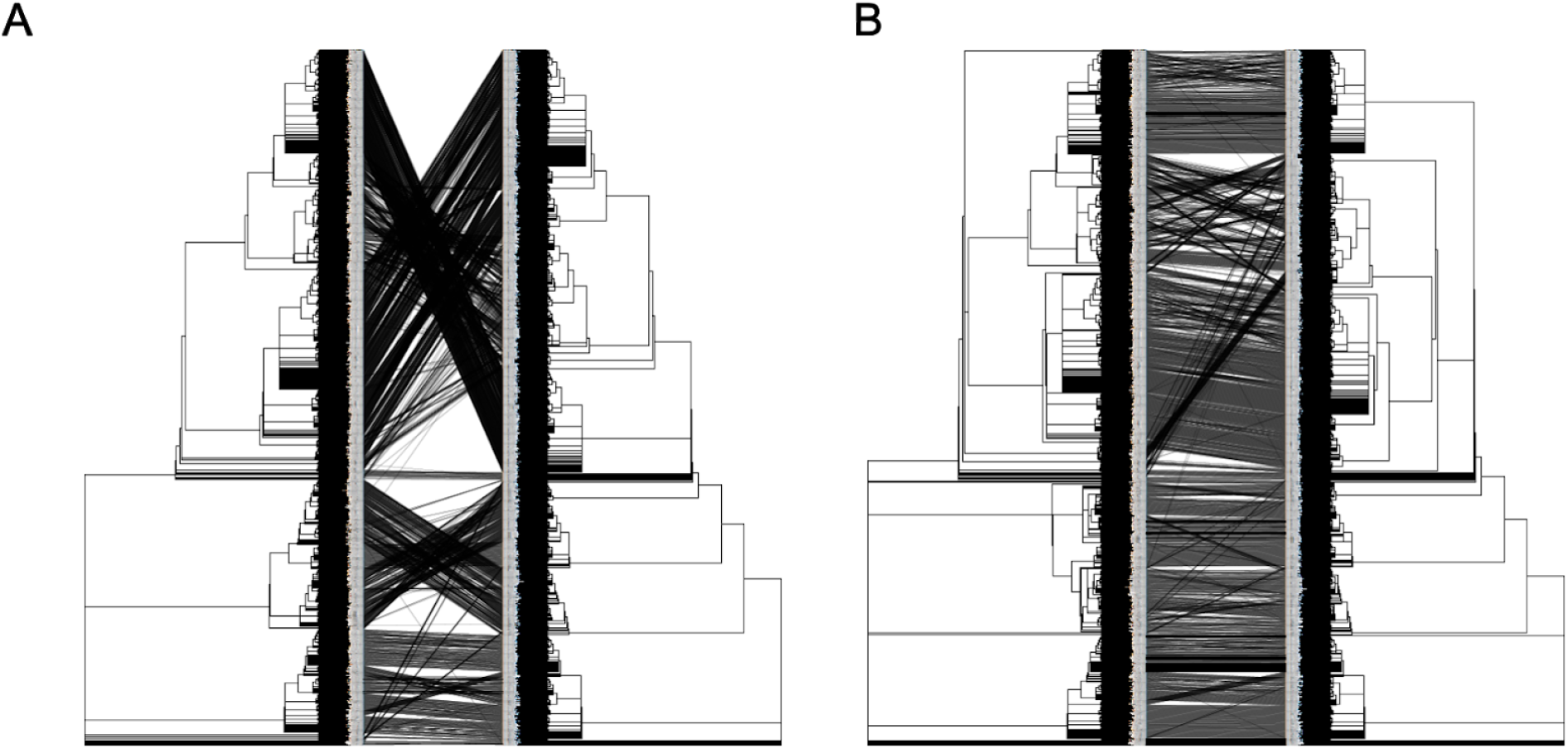
Tanglegrams for the two Nextstrain trees released on 04/19/2020 (left) and 04/20/2020 (right). (**A**) Without tree rotation, the tanglegram has a large mesh of connecting lines, making it hard to see the tree correspondence. (**B**) With trees rotated using RotTrees, the tanglegram is more visually appealing and the tree correspondence is a lot clearer.

**Figure S6:**
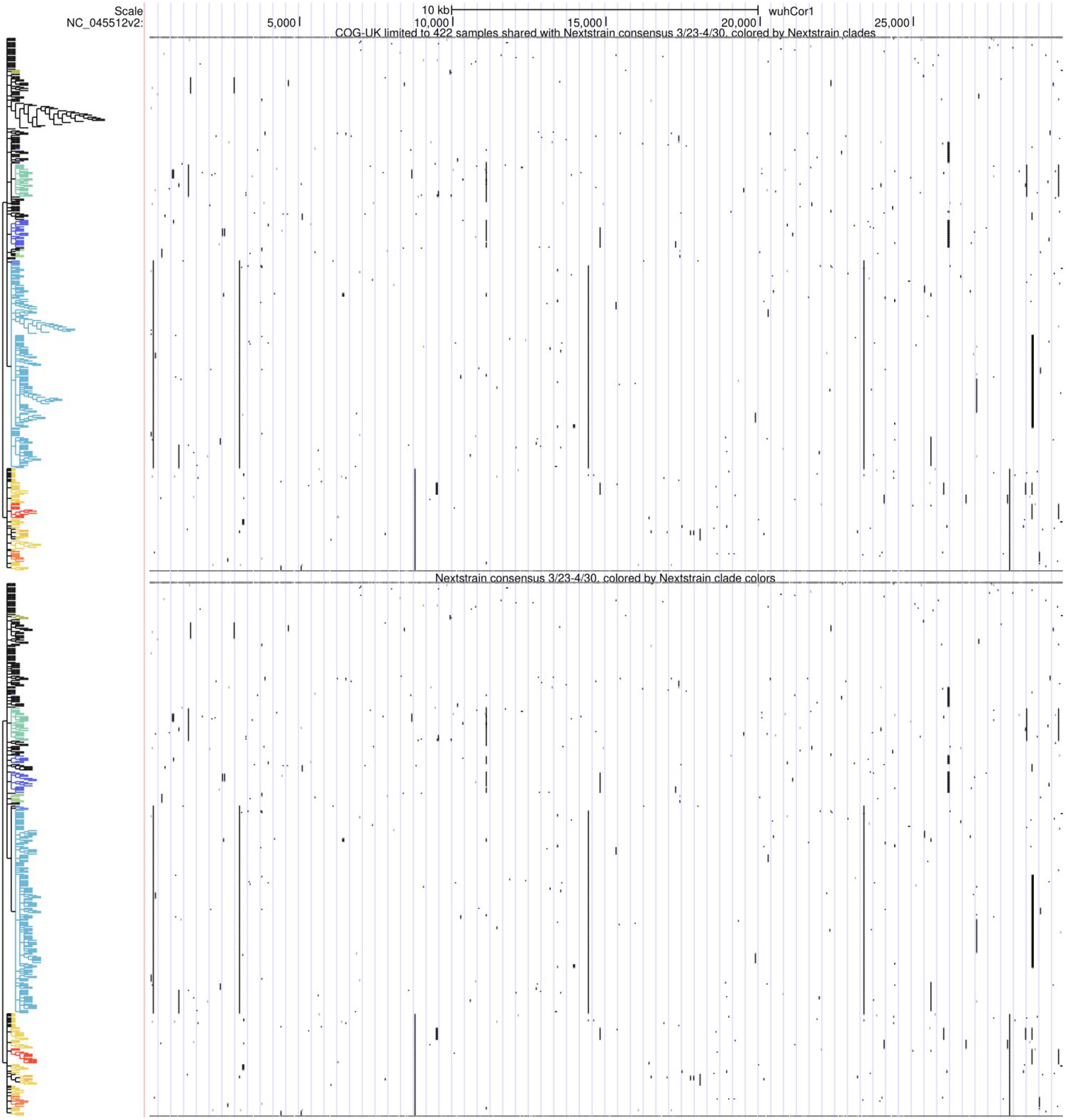
UCSC Genome Browser display of the trees from Figure 11C (COG-UK tree from 4/24, restricted to 422 samples in common with consensus tree of Nextstrain trees 3/23-4/30, and Nextstrain consensus tree), colored by Nextstrain clade assigned to sample. Interactive view: http://genome.ucsc.edu/s/SARS_CoV2/cogVsNsCladeColors

**Table S1.** Lab-associated mutations discovered in our dataset.

| SITE | Annotation | ARCTIC<br>Primer w/in<br>10bp | Alt allele<br>count | Parsim<br>ony | Explanation | Alternat<br>e Allele<br>Freque<br>ncy | Automa<br>ted Bin<br>Result | Numb<br>er of<br>Clade<br>s | Number of<br>lineages in<br>each<br>subclade (a,<br>b, c...) |
| --- | --- | --- | --- | --- | --- | --- | --- | --- | --- |
| G3564T | AACHANGE=<br>ORF1a:G1100<br>V |  | 14 | 7 | 92.86% of alternate allele<br>calls stem from<br>Microbiological Diagnostic<br>Unit Public Health<br>Laboratory | 0.0030<br>82 | highly<br>suspect | 7 | 1,1,2,2,2,5,1 |
| G8790T | AACHANGE=<br>ORF1a:G2842<br>V |  | 4 | 4 | 100.0% of alternate allele<br>calls stem from<br>Microbiological Diagnostic<br>Unit Public Health<br>Laboratory | 0.0008<br>81 | highly<br>suspect | 4 | 1,1,1,1 |
| G24933T | AACHANGE=S<br>:G1124V |  | 11 | 6 | 100.0% of alternate allele<br>calls stem from<br>Microbiological Diagnostic<br>Unit Public Health<br>Laboratory | 0.0024<br>22 | highly<br>suspect | 6 | 1,1,1,2,5,1 |
| G2198A | AACHANGE=<br>ORF1a:G645S | 8_left | 5 | 4 | 100.0% of alternate allele<br>calls stem from Erasmus<br>Medical Center | 0.0011<br>01 | highly<br>suspect | 4 | 2,1,1,1 |
| G3145T | AACHANGE=<br>ORF1a:L960F | 11_left | 9 | 6 | 100.0% of alternate allele<br>calls stem from Erasmus<br>Medical Center | 0.0019<br>82 | highly<br>suspect | 5 | 1,1,1,1,5 |
| A3778G |  | 13_left | 7 | 6 | 85.71% of alternate allele<br>calls stem from Erasmus<br>Medical Center | 0.0015<br>41 | highly<br>suspect | 6 | 1,2,1,1,1,1 |
| C6255T | AACHANGE=<br>ORF1a:A1997<br>V | 20_right | 12 | 9 | 83.33% of alternate allele<br>calls stem from Erasmus<br>Medical Center | 0.0026<br>42 | highly<br>suspect | 8 | 1,1,5,1,1,1,1<br>,1 |
| A4050C | AACHANGE=<br>ORF1a:N1262<br>T | 14_left_alt<br>4 | 18 | 11 | 100.0% of alternate allele<br>calls stem from KU Leuven,<br>Clinical and Epidemiological<br>Virology | 0.0039<br>63 | highly<br>suspect | 10 | 1,1,1,1,1,1,1<br>,1,1,9 |
| T8022G | AACHANGE=<br>ORF1a:V2586<br>G | 26_right | 5 | 5 | 100.0% of alternate allele<br>calls stem from KU Leuven,<br>Clinical and Epidemiological<br>Virology | 0.0011<br>01 | highly<br>suspect | 5 | 1,1,1,1,1 |
| T13402G | AACHANGE=<br>ORF1a:Y4379* | 44_right | 53 | 16 | 92.45% of alternate allele<br>calls stem from KU Leuven,<br>Clinical and Epidemiological<br>Virology | 0.0116<br>69 | highly<br>suspect | 12 | 1,5,1,24,2,1,<br>2,1,1,9,1,5 |
| A13947T |  | 47_left | 12 | 5 | 83.33% of alternate allele<br>calls stem from KU Leuven,<br>Clinical and Epidemiological<br>Virology | 0.0026<br>42 | highly<br>suspect | 6 | 1,2,5,2,1,1 |
| A24389C | AACHANGE=S<br>:S943P | 81_left | 23 | 5 | 95.65% of alternate allele<br>calls stem from KU Leuven,<br>Clinical and Epidemiological<br>Virology | 0.0050<br>64 | highly<br>suspect | 5 | 1,19,1,1,1 |
| G24390C | AACHANGE=S<br>:S943P | 81_left | 24 | 6 | 95.83% of alternate allele<br>calls stem from KU Leuven,<br>Clinical and Epidemiological<br>Virology | 0.0052<br>84 | highly<br>suspect | 6 | 1,1,19,1,1,1 |
| G1149T | AACHANGE=<br>ORF1a:G295V |  | 13 | 4 | 92.31% of alternate allele<br>calls stem from Department<br>of Health Technology and<br>Informatics, Faculty of<br>Health and Social Science,<br>The Hong Kong Polytechnic<br>University | 0.0028<br>62 | highly<br>suspect | 4 | 1,1,10,1 |
| C22802G | AACHANGE=S<br>:Q414E | 76_left,<br>76_left_alt<br>3 | 8 | 6 | 100.0% of alternate allele<br>calls stem from TGen North | 0.0017<br>61 | highly<br>suspect | 6 | 1,1,1,1,1,3 |
| T153G |  |  | 4 | 4 | 100.0% of alternate allele<br>calls stem from UW<br>Virology Lab | 0.0008<br>81 | highly<br>suspect | 4 | 1,1,1,1 |

**Table S2.**
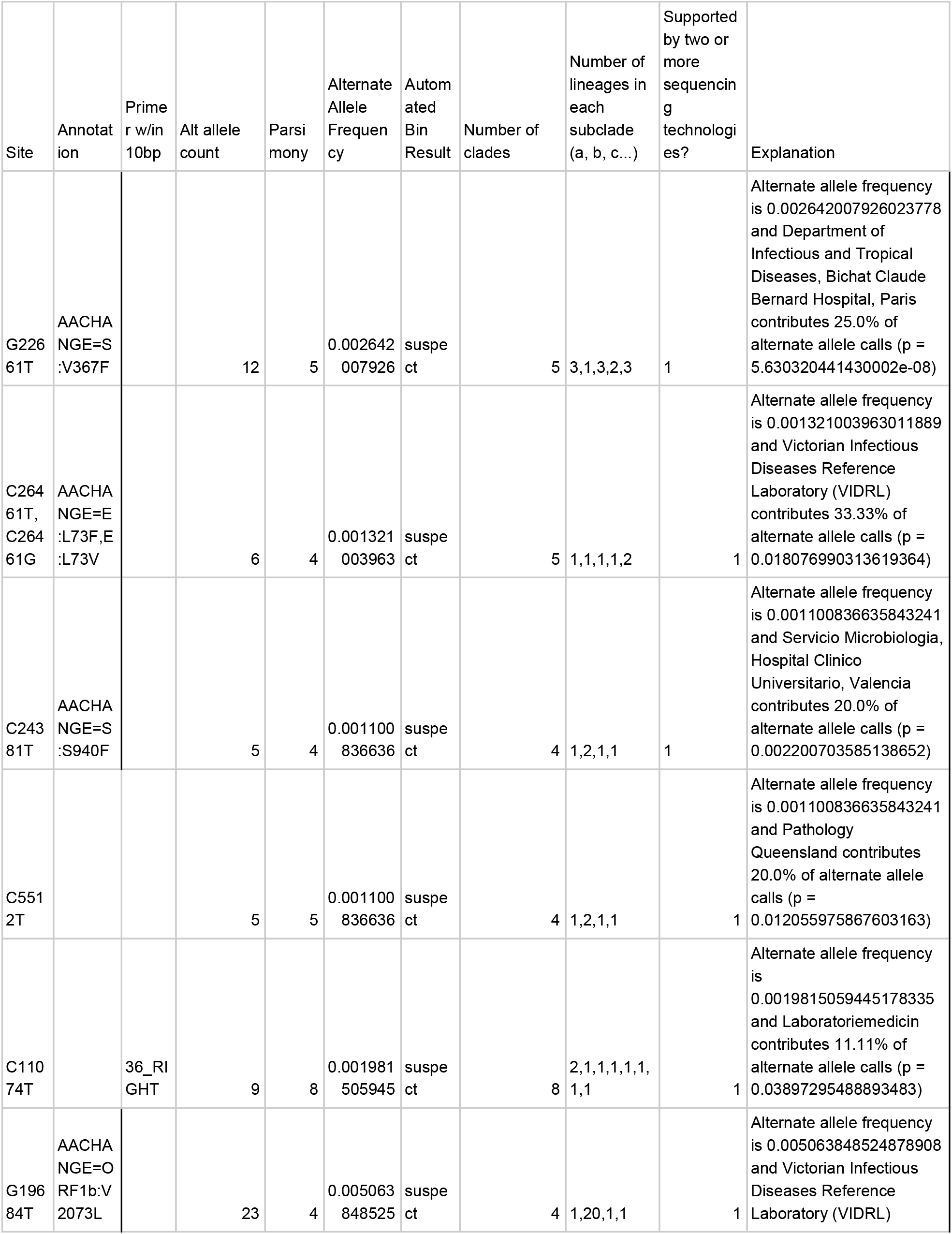
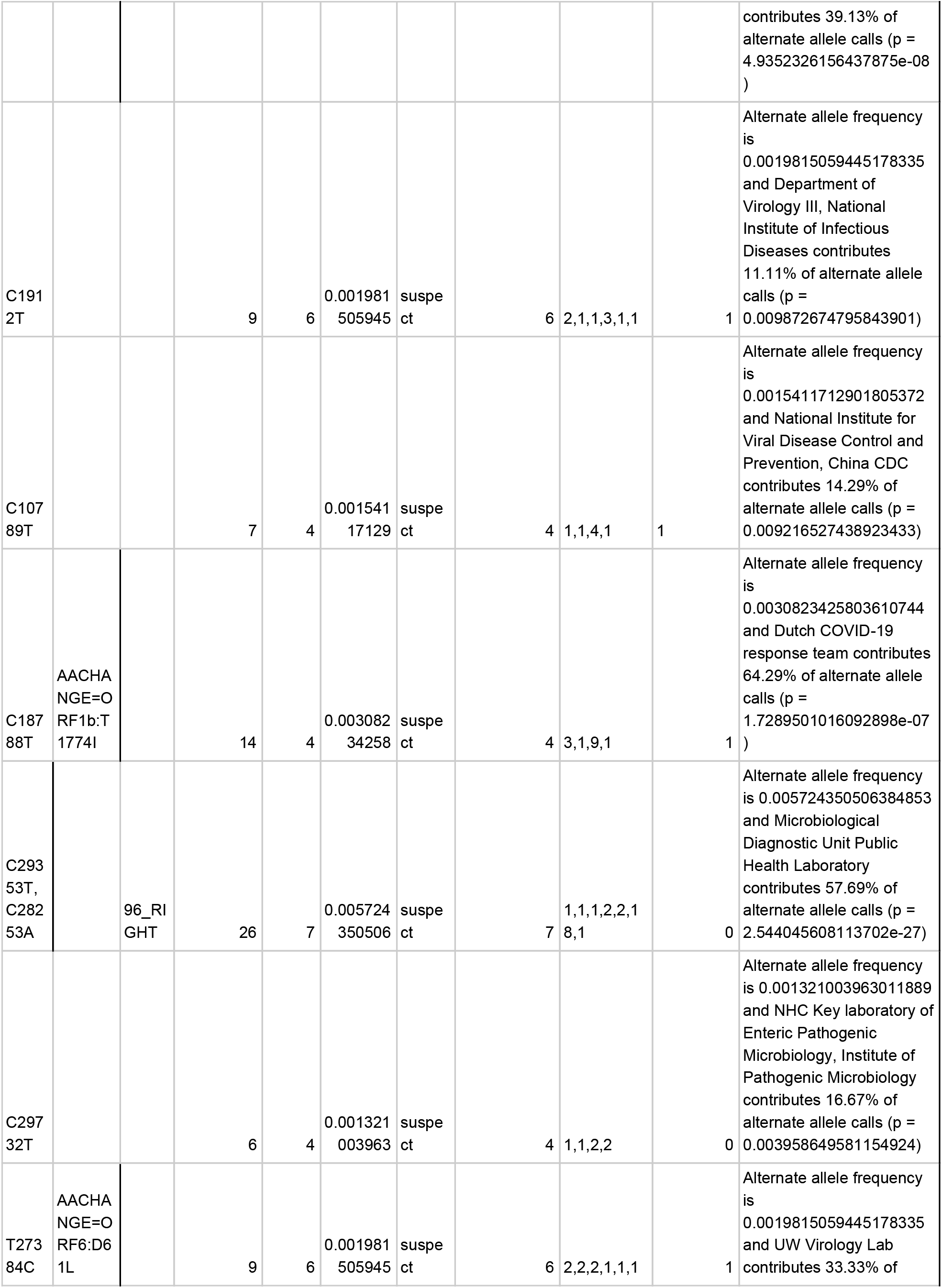
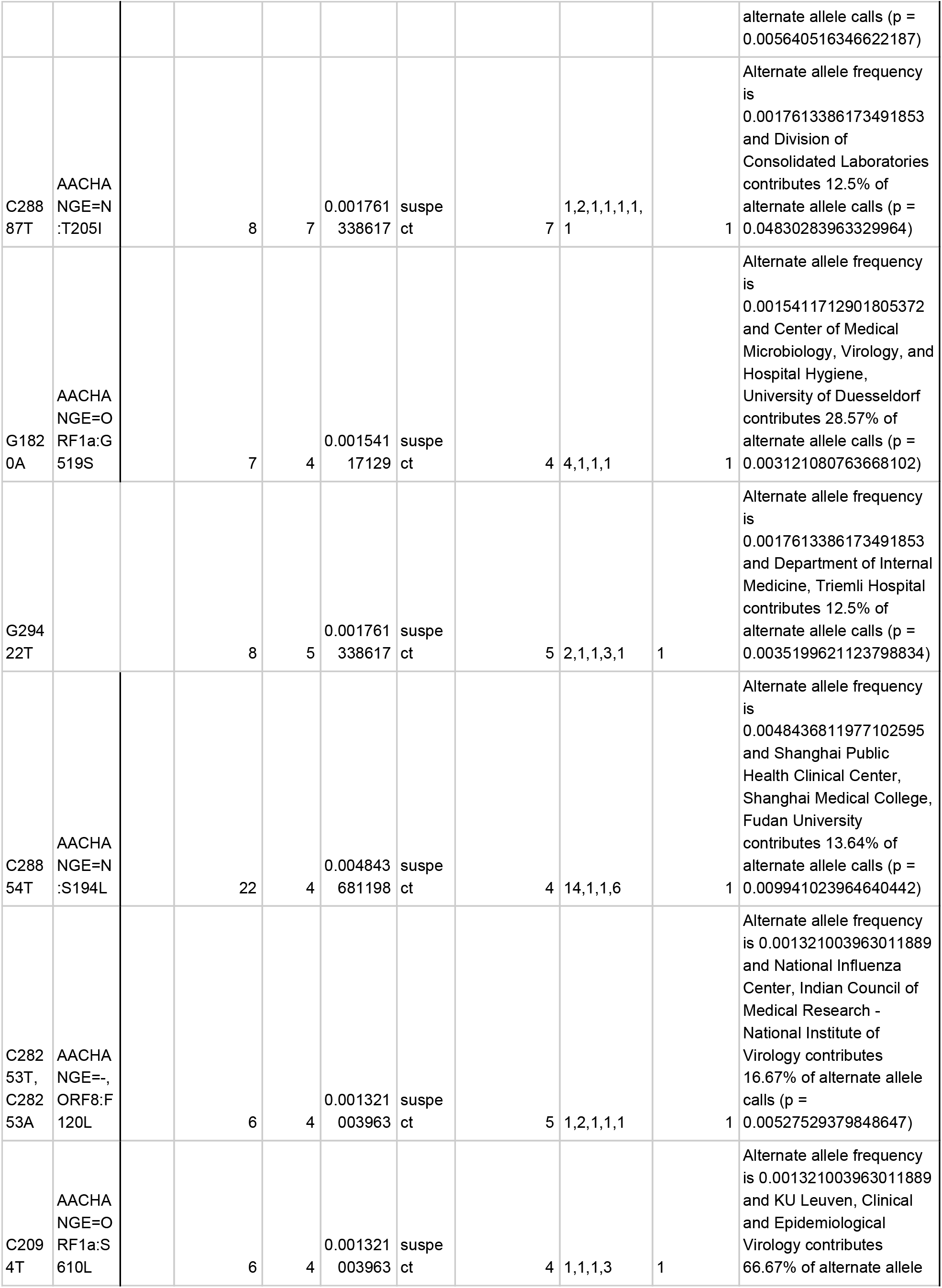
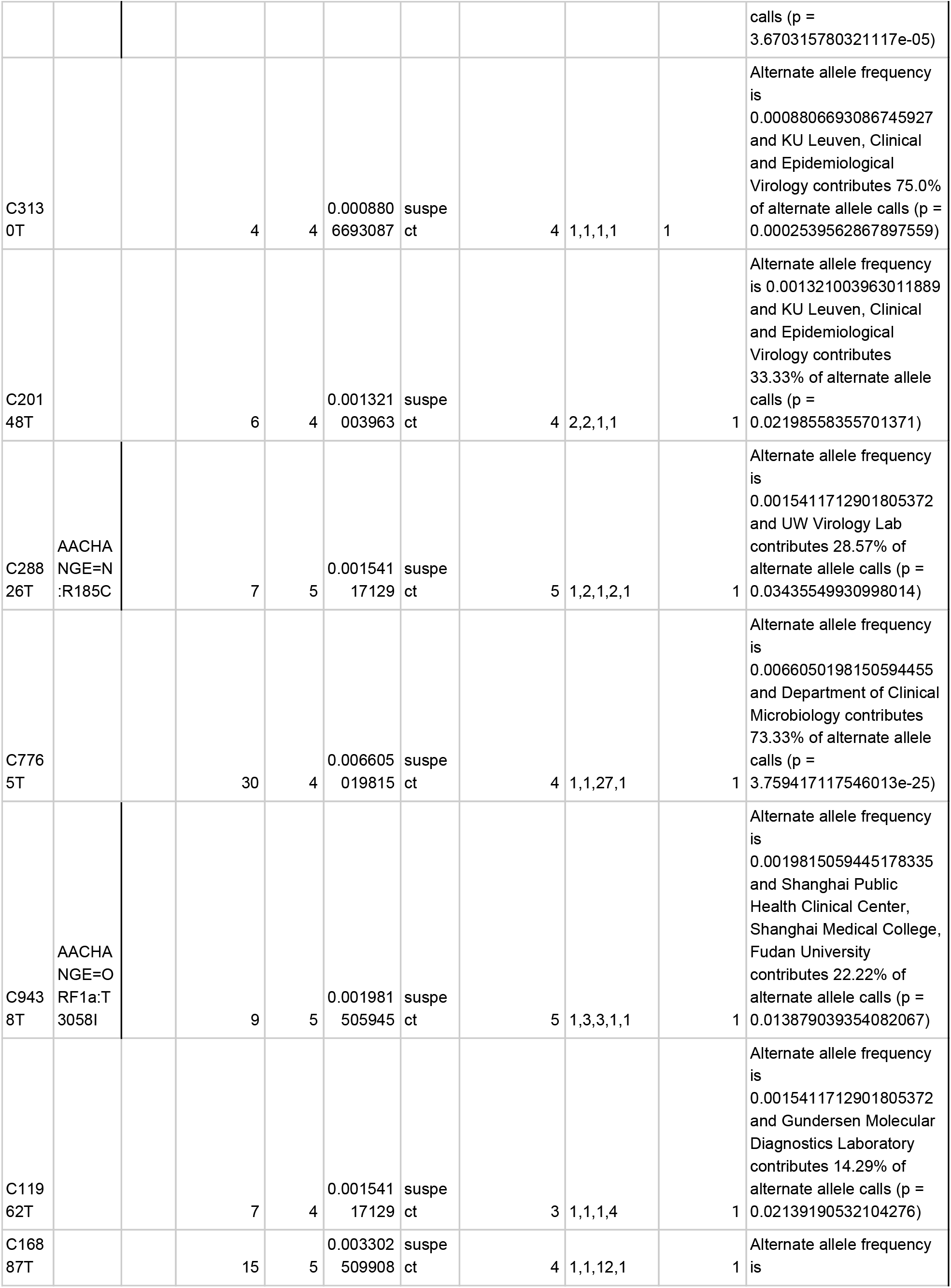
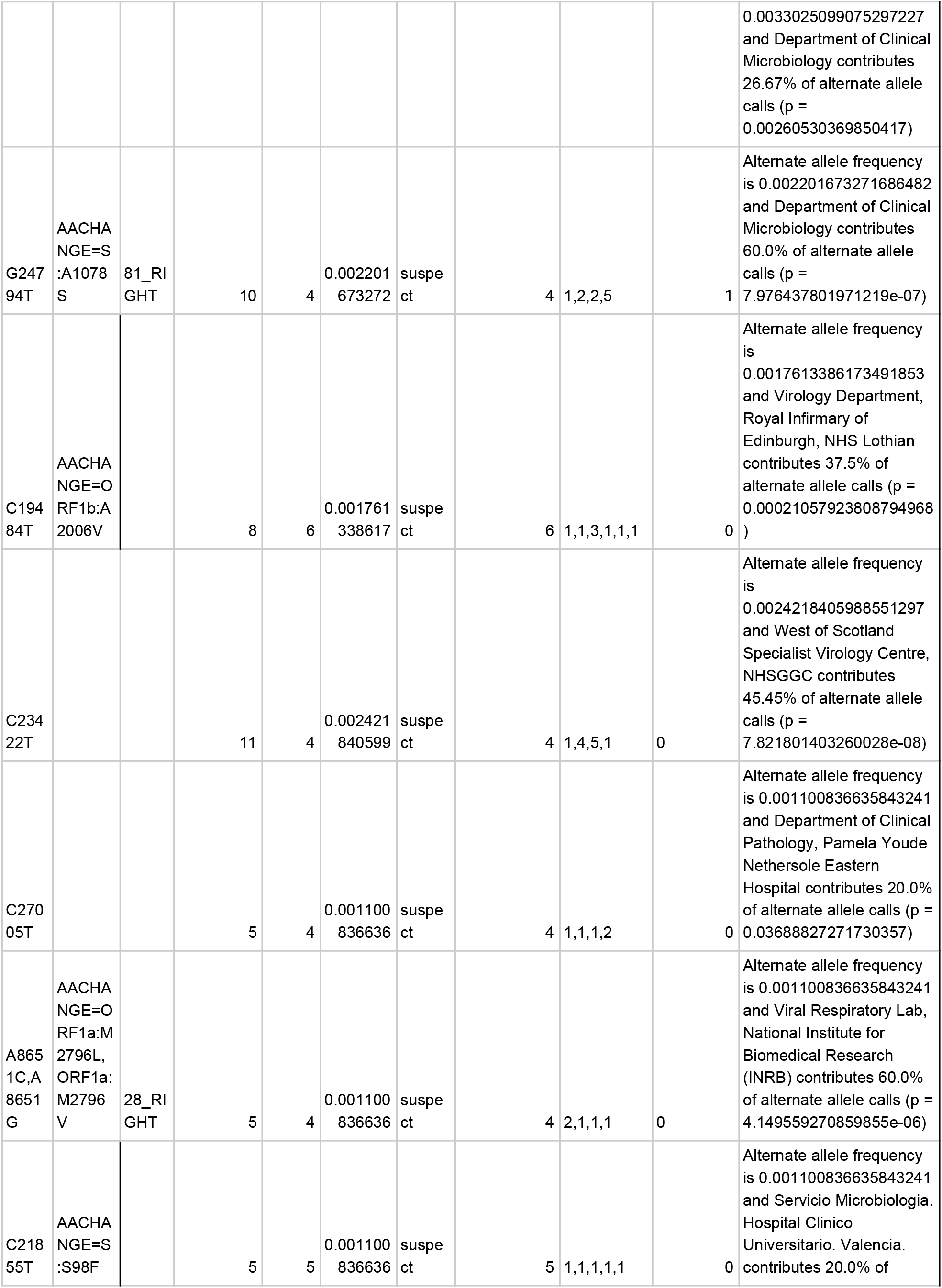
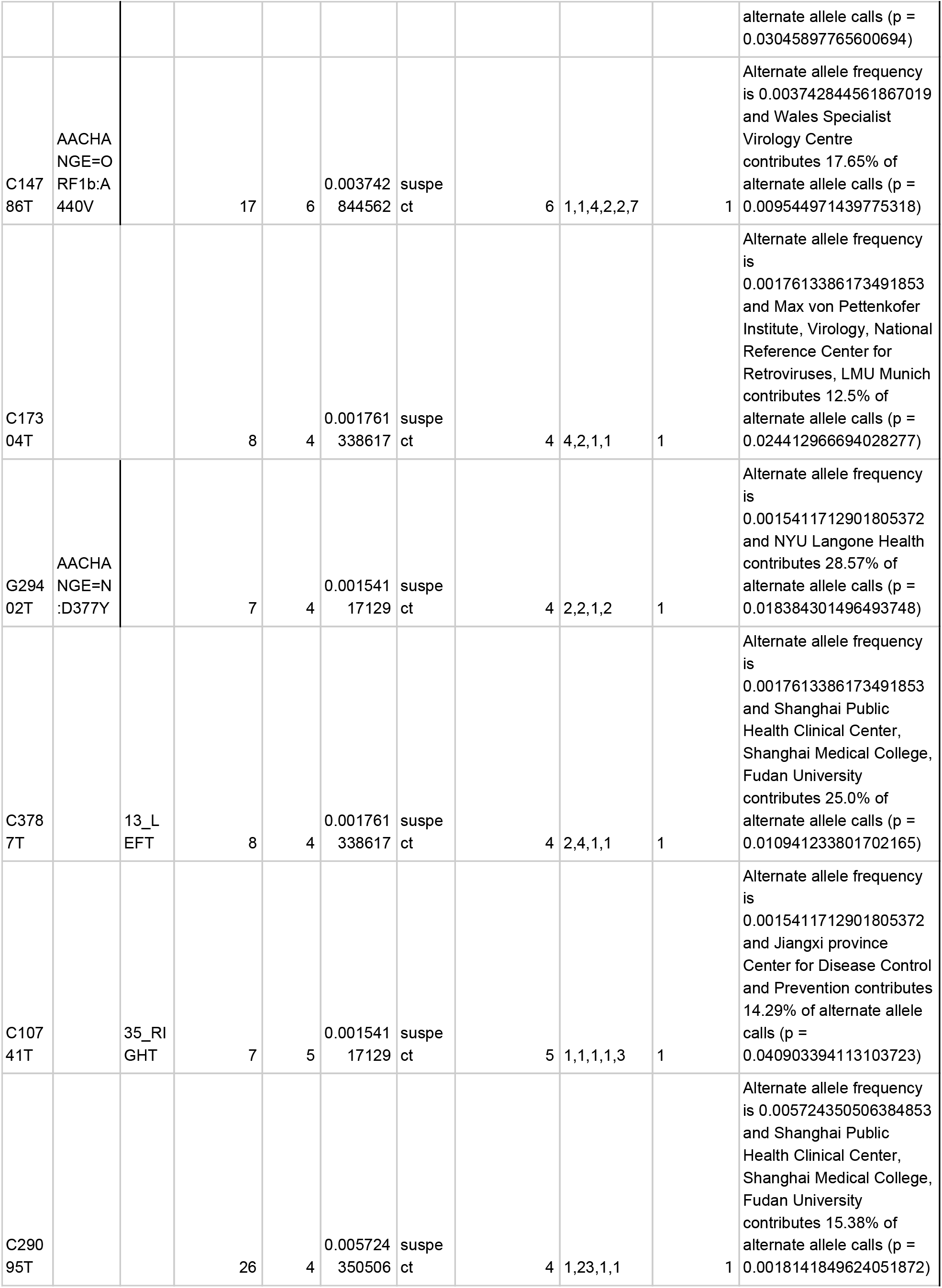
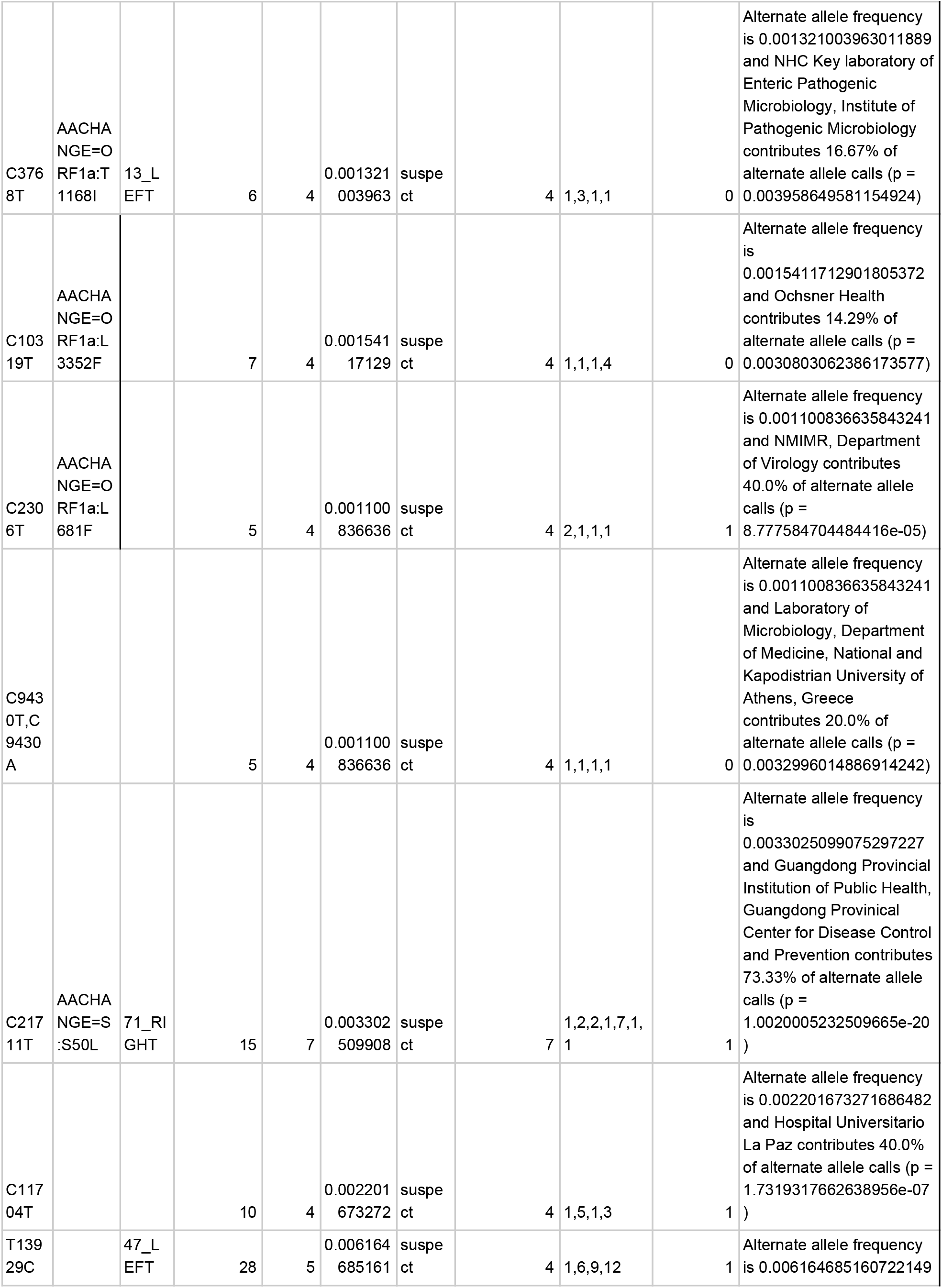
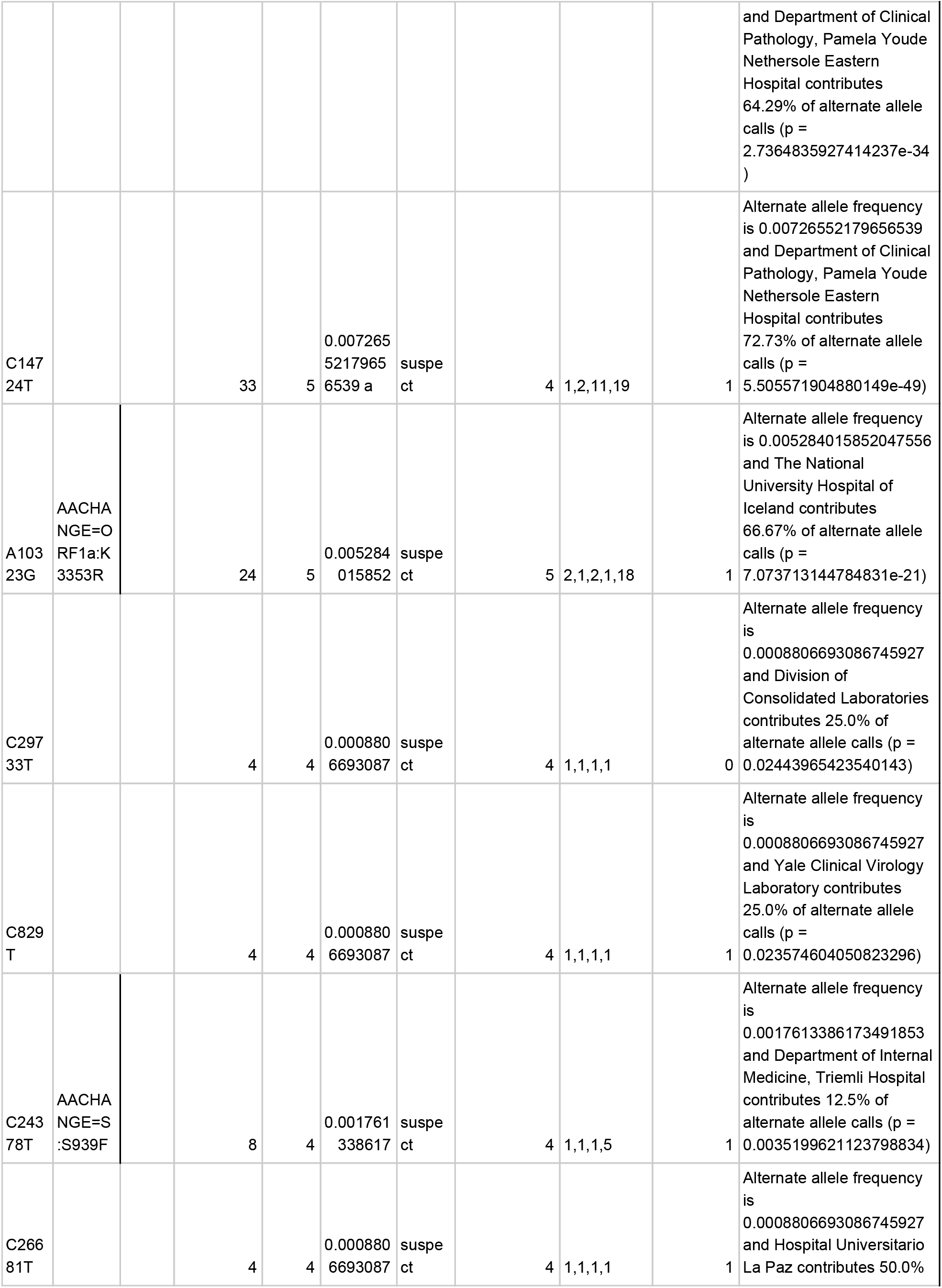
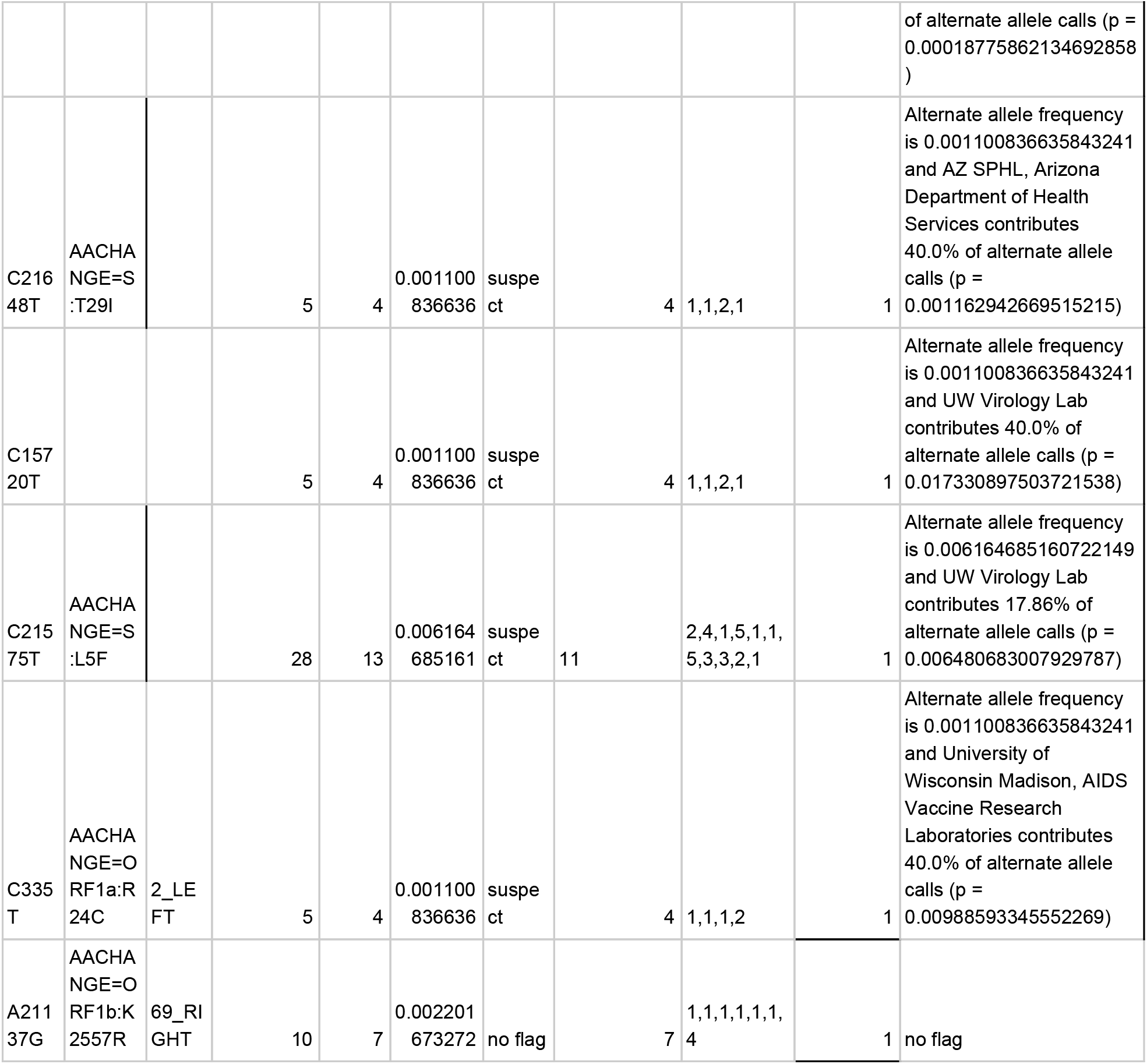
Highly recurrent, low alternate allele frequency mutations.

**Table S3.** High alternate allele frequency, highly recurrent sites.

| SITE | Annotation | ARTIC Primer<br>w/in 10bp | Alt allele count | Parsimony | Alternate Allele<br>Frequency | Automated Bin<br>Result |
| --- | --- | --- | --- | --- | --- | --- |
| C241T |  |  | 2793 | 7 | 0.6149273448 | no flag |
| C1059T | AACHANGE=OR<br>F1a:T265I |  | 834 | 7 | 0.1836195509 | no flag |
| C3037T,C3037A | AACHANGE=-,O<br>RF1a:F924L |  | 2793 | 7 | 0.6149273448 | no flag |
| C8782T |  |  | 673 | 4 | 0.1481726112 | no flag |
| G11083T | AACHANGE=OR<br>F1a:L3606F | 36_RIGHT | 504 | 26 | 0.1109643329 | no flag |
| C14408T | AACHANGE=OR<br>F1b:P314L |  | 2763 | 9 | 0.608322325 | no flag |
| C14805T |  |  | 397 | 5 | 0.08740642889 | no flag |
| C15324T |  |  | 238 | 12 | 0.05239982387 | no flag |
| C18060T |  | 59_RIGHT | 296 | 5 | 0.06516952884 | no flag |
| C18877T |  |  | 133 | 4 | 0.02928225451 | no flag |
| A20268G |  |  | 154 | 6 | 0.03390576838 | no flag |
| A23403G | AACHANGE=S:<br>D614G |  | 2792 | 7 | 0.6147071775 | no flag |
| C24034T |  |  | 67 | 4 | 0.01475121092 | no flag |
| G25563T | AACHANGE=OR<br>F3a:Q57H |  | 1082 | 8 | 0.238221048 | no flag |
| G26144T | AACHANGE=OR<br>F3a:G251V |  | 391 | 4 | 0.08608542492 | no flag |
| C27046T | AACHANGE=M:<br>T175M |  | 136 | 7 | 0.02994275649 | no flag |
| C28311T | AACHANGE=N:<br>P13L |  | 59 | 4 | 0.0129898723 | no flag |
| A29700G |  | 97_RIGHT | 49 | 6 | 0.01078819903 | no flag |

**Table S4.** Lab-associated, low parsimony score sites.

| SITE | Annotation | ARTIC Primer w/in 10bp | Alt allele count | Parsimony | Explanation | Alternate Allele Frequency | Number of Clades | Number of lineages in each subclade (a, b, c...) |
| --- | --- | --- | --- | --- | --- | --- | --- | --- |
| T7438C | AACHAN<br>GE=ORF<br>1a:Y4379<br>* |  | 18 | 1 | 100.0% of alternate allele calls stem from Microbiological Diagnostic Unit Public Health Laboratory | 0.0039630 | 1 | 18 |
| A26864G |  | 89_LEFT,89_LEFT_alt2 | 13 | 1 | 100.0% of alternate allele calls stem from Microbiological Diagnostic Unit Public Health Laboratory | 0.0299428 | 1 | 13 |
| T1570C |  | 6_LEFT | 13 | 1 | 100.0% of alternate allele calls stem from Erasmus Medical Center | 0.0028622 | 1 | 13 |
| A11438G | AACHAN<br>GE=ORF<br>1a:N3725<br>D |  | 10 | 1 | 100.0% of alternate allele calls stem from Erasmus Medical Center | 0.0022017 | 1 | 10 |
| G12550A |  |  | 10 | 1 | 100.0% of alternate allele calls stem from Erasmus Medical Center | 0.0022017 | 1 | 10 |
| C16762T | AACHAN<br>GE=ORF<br>1b:L1099<br>F | 56_LEFT | 16 | 2 | 81.25% of alternate allele calls stem from Erasmus Medical Center | 0.0035227 | 2 | 3,13 |
| C20740A | AACHAN<br>GE=ORF<br>1b:Q2425<br>K |  | 10 | 1 | 100.0% of alternate allele calls stem from Erasmus Medical Center | 0.0030823 | 1 | 10 |
| C21590T |  |  | 14 | 1 | 85.71% of alternate allele calls stem from GIGA Medical Genomics | 0.0030823 | 1 | 14 |
| C9962T | AACHAN<br>GE=ORF<br>1a:H3233<br>Y |  | 33 | 2 | 87.88% of alternate allele calls stem from Department of Health Technology and Informatics, Faculty of Health and Social Science, The Hong Kong Polytechnic University | 0.0072655 | 2 | 1,32 |
| T215<br>84G | AACHAN<br>GE=S:L8<br>V |  | 31 | 2 | 87.1% of alternate allele calls<br>stem from Department of Health<br>Technology and Informatics,<br>Faculty of Health and Social<br>Science, The Hong Kong<br>Polytechnic University | 0.0068252 | 2 | 13,18 |
| C514<br>2T | AACHAN<br>GE=ORF<br>1a:T1626I |  | 13 | 1 | 100.0% of alternate allele calls<br>stem from deCODE genetics | 0.0028622 | 1 | 13 |
| C240<br>54T | AACHAN<br>GE=S:A8<br>31V |  | 11 | 2 | 90.91% of alternate allele calls<br>stem from deCODE genetics | 0.0024218 | 2 | 1,10 |
| A259<br>58G |  |  | 10 | 1 | 100.0% of alternate allele calls<br>stem from deCODE genetics | 0.0022017 | 1 | 10 |
| C256<br>69T | AACHAN<br>GE=ORF<br>3a:H93Y | 84_RIGHT | 15 | 1 | 100.0% of alternate allele calls<br>stem from Public Health Wales<br>Microbiology Cardiff | 0.0033025 | 1 | 15 |
| G288<br>51T | AACHAN<br>GE=N:S1<br>93I |  | 34 | 2 | 97.06% of alternate allele calls<br>stem from Public Health Wales<br>Microbiology Cardiff | 0.0074857 | 2 | 1,33 |
| C267<br>50T |  |  | 17 | 1 | 100.0% of alternate allele calls<br>stem from WHO National<br>Influenza Centre Russian<br>Federation | 0.0037428 | 1 | 17 |
| T833<br>C | AACHAN<br>GE=ORF<br>1a:F190L |  | 34 | 1 | 82.35% of alternate allele calls<br>stem from UW Virology Lab | 0.0074857 | 1 | 34 |
| G295<br>53A |  |  | 22 | 1 | 95.45% of alternate allele calls<br>stem from UW Virology Lab | 0.0048437 | 1 | 22 |
| G387<br>1T | AACHAN<br>GE=ORF<br>1a:K1202<br>N |  | 16 | 2 | 93.75% of alternate allele calls<br>stem from University of<br>Wisconsin Madison, AIDS<br>Vaccine Research Laboratories | 0.0035227 | 2 | 1,15 |
| G114<br>17T | AACHAN<br>GE=ORF<br>1a:V3718<br>F |  | 31 | 1 | 93.55% of alternate allele calls<br>stem from University of<br>Wisconsin Madison, AIDS<br>Vaccine Research Laboratories | 0.0068252 | 1 | 31 |
| T140<br>73C |  |  | 28 | 1 | 100.0% of alternate allele calls<br>stem from University of<br>Wisconsin Madison, AIDS<br>Vaccine Research Laboratories | 0.0061647 | 1 | 28 |
| G157<br>60A | AACHAN<br>GE=ORF<br>1b:G765S |  | 11 | 2 | 81.82% of alternate allele calls<br>stem from University of<br>Wisconsin Madison, AIDS<br>Vaccine Research Laboratories | 0.0024218 | 2 | 1,10 |
| T178<br>77C |  |  | 23 | 1 | 100.0% of alternate allele calls<br>stem from University of<br>Wisconsin Madison, AIDS<br>Vaccine Research Laboratories | 0.0050638 | 1 | 23 |
| C203<br>16T |  |  | 23 | 1 | 100.0% of alternate allele calls<br>stem from University of<br>Wisconsin Madison, AIDS<br>Vaccine Research Laboratories | 0.0050638 | 1 | 23 |
| A239<br>47G |  |  | 28 | 1 | 100.0% of alternate allele calls<br>stem from University of<br>Wisconsin Madison, AIDS<br>Vaccine Research Laboratories | 0.0061647 | 1 | 28 |

**Table S5.**
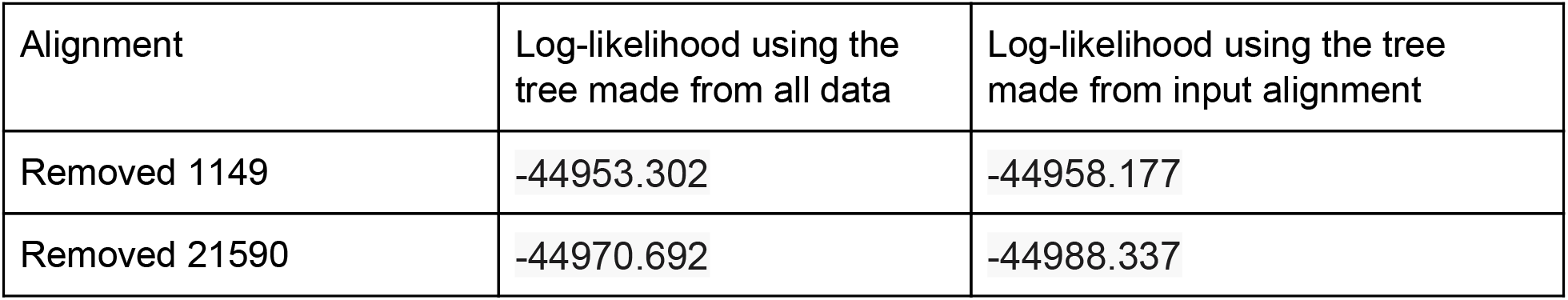
Log-likelihood of outlier alignments based on entropy-weighted total distance.

**Table S6:** GISAID IDs whose clade annotation changed in the Nextstrain tree from 4/19/2020 to 3/30/2020, and from 4/19/2020 to 4/28/2020.

| Clade change | GISAID IDs |  |
| --- | --- | --- |
|  | 04/19 -> 03/30 | 04/19 -> 04/28 |
| A2 -> A2a | 418278, 418281 | 418278, 418281, 420357,<br>420358, 420359, 420365,<br>420431, 420853, 421652,<br>422426 |
| B -> B1 | 404895, 408478, 411060,<br>413855, 413862, 415584, | 404895, 408478, 411060,<br>413862, 415584, 415588, |
|  | 415588, 415589, 416419,<br>416473, 416886 | 415589, 416473, 416886,<br>418822, 418825, 418826,<br>418829, 418830, 418840,<br>418843, 418848, 418850,<br>418852, 418854, 418993 |

